# Characterizing semantic compositions in the brain: A model-driven fMRI re-analysis

**DOI:** 10.1101/2025.04.25.650708

**Authors:** Marco Ciapparelli, Marco Marelli, William Graves, Carlo Reverberi

**Affiliations:** Department of Psychology, University of Milano-Bicocca, Milan, Italy; Milan Center for Neuroscience - NeuroMI, University of Milano-Bicocca, Milan, Italy; Department of Psychology, Rutgers University, Newark, NJ, 07102, USA

**Author notes:** **Corresponding author:** Marco Ciapparelli, Department of Psychology, University of Milano-Bicocca, Italy.

## Abstract

Semantic composition allows us to construct complex meanings (e.g., “dog house”, “house dog”) from simpler constituents (“dog”, “house”). So far, neuroimaging studies have mostly relied on high-level contrasts (e.g., meaningful > non-meaningful phrases) to identify brain regions sensitive to semantic composition. However, such an approach is less apt at addressing *how* composition is carried out, namely what functions best characterize the integration of constituent concepts. To address this limitation, we rely on simple computational models to explicitly characterize alternative compositional operations, and use representational similarity analysis to compare the representations of models to those of target regions of interest within the general semantic network. To better target composition beyond specific task demands, we re-analyze fMRI data aggregated from four published studies (N = 85), all employing two-word combinations but differing in task requirements. Converging evidence from confirmatory and exploratory analyses reveals compositional representations in the *pars triangularis* of the left inferior frontal gyrus (BA45), even when analyses are restricted to a subset where the task did not require participants to actively engage in semantic processing. These results suggest that BA45 represents combinatorial information automatically across task demands, and further characterize these combinatorial representations as resulting from the (symmetric) intersection of constituent features. Additionally, a cluster of compositional representations emerges in the left middle superior temporal sulcus, while semantic, but not compositional, representations are observed in the left angular gyrus. Overall, our work clarifies which brain regions represent semantic information compositionally across different contexts and task demands, and qualifies which operations best describe composition.

## 1. Introduction

### 1.1 The neural bases of semantic composition

One of the core features of human cognition is the ability to construct complex meanings from simpler constituents. For example, the meaning of the complex word “party boat” – a boat that is used by people to celebrate – can be understood by combining the constituent concepts of “boat” and “party”, even if you have never heard of it. These “minimal combinations”, namely pairs of content words (e.g., noun-noun compound words), offer cognitive scientists a privileged window into conceptual combination: rather than producing sentences or coining new words, minimal phrases rely on the ability of people to infer the implicit meaning that links constituents. By looking at how minimal combinations are represented and processed, cognitive scientists compare and refine theories of semantic composition. For example, some theories propose that combinations are formed by selecting an implicit relational interpretation among a set of competitors (Gagné & Shoben, 1997), while others (Wisniewski, 1996) posit a qualitative distinction between combinations based on attribute modification (e.g., “robin hawk” is “a hawk with a red underbelly”) and those based on thematic relations (e.g., “taxi driver” is “a driver of taxis”). In this context, neuroimaging evidence can help adjudicate some theories by tapping into unobservable representations (Mack et al., 2013). Indeed, minimal combinations have been the object of many neuroimaging studies (Coutanche et al., 2018; Frankland & Greene, 2020a), which have largely taken a univariate approach focused on isolating brain activity related to semantic composition. More specifically, these approaches leverage *a-priori-*defined, high-level dimensions of interest (e.g., plausibility, familiarity, literalness), which, through appropriate contrasts (e.g., simple vs. complex words, familiar vs. novel complex words), are used to predict changes in the single-voxel signal.

Following this approach, neuroimaging studies have identified multiple regional and network specializations sensitive to factors attributable to conceptual combination. Of those, four regions of interest – from now on, *core ROIs* – have been consistently observed (Coutanche et al., 2018): the left anterior temporal lobe (LATL), the left and right angular gyri (LAG, RAG), and the left inferior frontal gyrus (LIFG). In line with its role as a multimodal semantic hub (Lambon Ralph et al., 2010; Ralph et al., 2016), the LATL appears to support the integration of semantic information to specify a single entity (Coutanche et al., 2018; Frankland & Greene, 2020a), both at the level of features (e.g., “green”, “round”, and “tart” as features of the concept “lime”; Coutanche & Thompson-Schill, 2015) and the level of concepts (e.g., the modifier “red” concept specifying the head concept “boat” in the compound “red boat”; Bemis & Pylkkänen, 2011). Consistent with this, evidence from MEG studies suggests that early LATL activation is modulated by how much a concept is made specific by the combination (Pylkkänen, 2019; Westerlund & Pylkkänen, 2014; Zhang & Pylkkänen, 2015; Ziegler & Pylkkänen, 2016), and that the LATL supports combinations based on extracting shared features, or *intersective conjunctions* (e.g., “The girls are tall and blonde”, with both “tall” and “blonde” referring to the same entity; Poortman & Pylkkänen, 2016), with fMRI studies advancing analogous considerations (Baron & Osherson, 2011; Coutanche & Thompson-schill, 2015). Finally, semantic combination in LATL has been characterized as largely independent of plausibility and syntactic composition (even though these factors might lead to additive effects; Parrish & Pylkkänen, 2022). Instead, RAG and, most notably, LAG activation have been related to compound meaningfulness (Forgács et al., 2012; Graessner et al., 2021; Graves et al., 2010; Price et al., 2015; Price et al., 2016). Concerning AG computations, Coutanche et al., (2018) argue that AG supports *relational combinations* (see Boylan et al., 2017), namely those mediated by an implicit thematic relation linking constituents (e.g., “crayon box” is “a box that contains crayons”), consistent with evidence showing that AG represents thematic relations more broadly (Wang et al., 2020; Zhang et al., 2022; Schwartz et al., 2011). However, Ralph et al., (2016) and Humphreys et al., (2021) raised the possibility that, in line with its contribution to the default mode network, AG activation may simply index task difficulty (but see Boylan et al., 2017). Finally, LIFG (specifically *pars triangularis* and *pars orbitalis*) involvement in semantic composition studies has been linked to *decreased* compound meaningfulness (Forgács et al., 2012; Graessner, Zaccarella, Friederici, et al., 2021; Graves et al., 2010; Molinaro et al., 2015) and increased feature uncertainty (Solomon & Thompson-Schill, 2020). It has therefore been suggested that the involvement of this region could be explained by the greater semantic control demands (e.g., selection of the appropriate features to integrate; Coutanche et al., 2018) imposed by less meaningful combinations (Graves et al., 2010, Molinaro et al., 2015), consistent with LIFG role within the semantic control network (Jackson, 2021). Besides these core ROIs, other regions have been implicated in semantic composition, most notably the ventromedial prefrontal cortex (vmPFC), posterior cingulate cortex (PCC), and left middle-superior temporal sulcus (lmSTS) (Frankland & Greene, 2020a; Pylkkänen, 2019), which, together with the LATL, AG, and LIFG, mostly lie within the broader semantic network (Binder et al., 2009; Jackson, 2021) and the default mode network (Frankland & Greene, 2020a).

While these studies have provided a rich picture of the neural bases of semantic composition, their approach based on high-level contrast (e.g., interpretable vs. uninterpretable, literal vs. figurative combinations) is more apt at isolating composition-related brain activity, rather than characterizing the transformations that define composition itself. For example, contrasting plausible and non-plausible combinations might reveal which regions engage in semantic composition and are thus modulated by the degree to which constituents are combinable. However, such results do not tell us how constituent meanings are transformed and integrated (e.g., by feature intersection or “intersective conjunction”, by attribution or subordination; Scalise & Bisetto, 2005). In other words, while this approach can reveal whether and where composition is accomplished, it does not shed light on *how* the combination is carried out. We do not claim that high-level contrasts cannot be of help: indeed, one can devise contrasts that target specific composition types (e.g., attributive vs. relational combinations; Boylan et al., 2017). However, this approach is still limited in a number of ways. First, it relies on high-level distinctions that might not adequately describe the mechanisms of combination. For example, Marelli et al., (2017) showed that some effects of conceptual combination can be accounted for by a general mechanism based on extracting statistical regularities over (compound) word usage. In this context, distinctions among combination types are better described as the by-product of a general distributional system and are thus less amenable to be pigeonholed into discrete contrasts. Second, many combinations are amenable to more than one interpretation (Schmidtke et al., 2018) and composition type (Kenett & Thompson-Schill, 2020). Perhaps most notably, the meaning of familiar combinations (e.g., “swordfish”) can be both retrieved from long-term memory (like single words) or can be generated by an active process of combination (like sentences), with theoretical and empirical reasons suggesting that the two are attempted simultaneously (Libben, 2014). In other words, the same combinations can be processed and thus represented in alternative ways, both compositional and non-compositional, which the brain might carry out in parallel (Baggio, 2021). Studying these distinctions with contrasts would thus be difficult or even impossible, since the same stimulus may belong to different contrast levels depending on specific conditions (e.g., context, task demands, or individual preferences), or even irrespective of conditions (e.g., if both retrieval and combinations were attempted in parallel and automatically; Libben, 2014). We argue that these limitations can be addressed by relating brain activity to computational models of semantic combination. Indeed, by formally defining both semantic representations and the transformations that operate over them, the explanatory performance of computational models can be directly related to the specific type of semantic composition they implement.

### 1.2 Semantic composition from a distributional semantics perspective

Based on the hypothesis that words with similar meanings tend to occur in similar contexts (Harris, 1954), distributional semantics models (DSMs) exploit the co-occurrence statistics of large collections of text to generate semantic spaces, namely high-dimensional vector spaces where word meanings are represented as vectors (i.e., points in a semantic space) whose distance is inversely proportional to their semantic similarity (Figure 1). DSMs have shown good performance on diverse tasks (Baroni, Dinu, et al., 2014), and are considered cognitive models of semantic memory (Günther et al., 2019; Kumar, 2020). Importantly for our purpose, many studies have demonstrated representational similarities between these models and fine-grained activation patterns in the brain (Carota et al., 2017; Mitchell et al., 2008; Pereira et al., 2018; Zhang et al., 2020).

**Figure 1:**
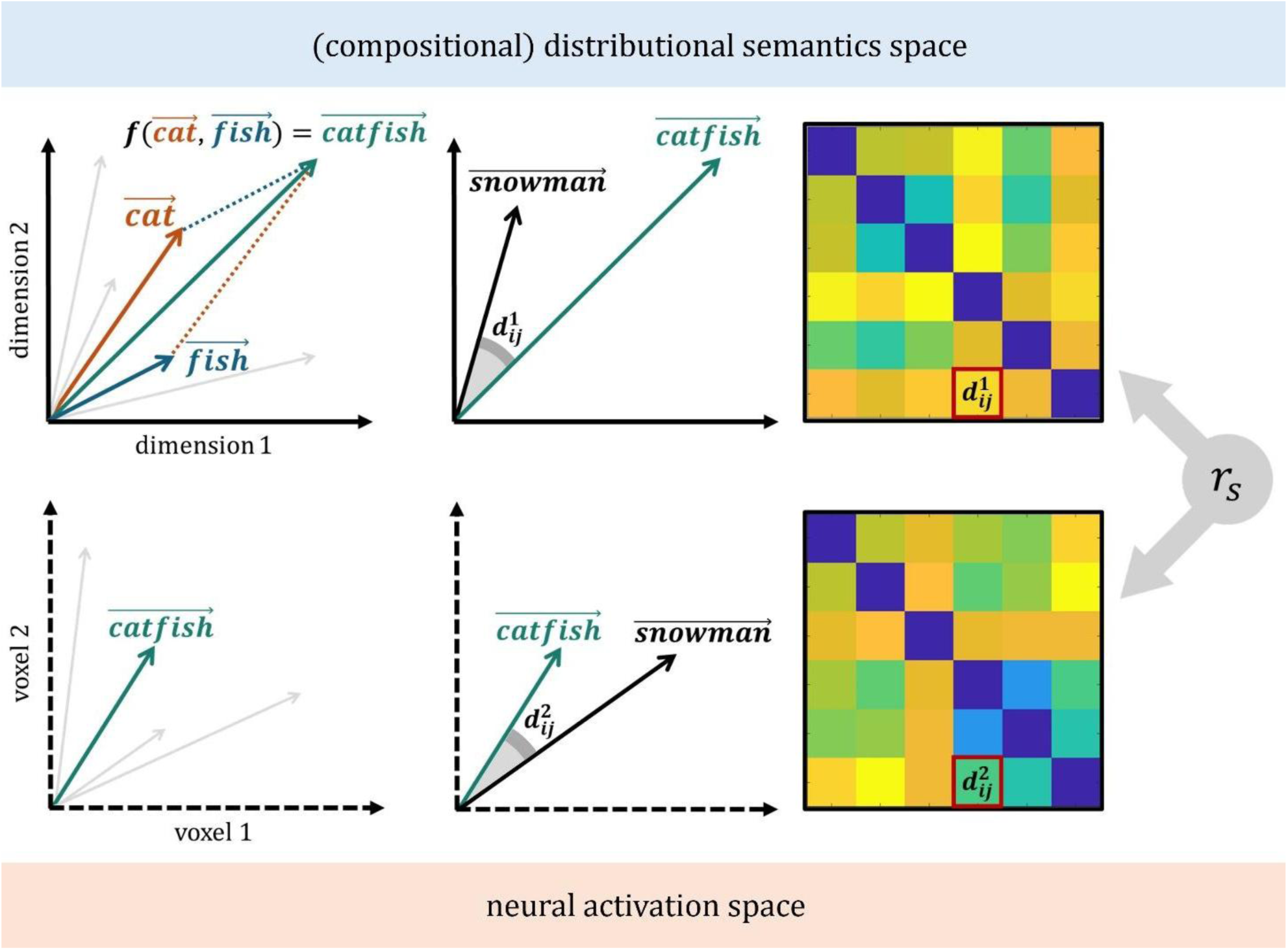
Representational similarity analysis (RSA) between the space of a cDSM (top) and the space of an ROI (bottom). The top left image depicts a simplified semantic space where the meaning of “cat” and “fish” are represented by 2-dimensional vectors. A cDSM, namely the function *f*, takes the two vectors as argument to generate the vector of their combination, “catfish.” Then, the semantic dissimilarity of “catfish” and another combination, “snowman”, is quantified. The dissimilarity 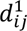 between combinations *i* and *j* defines the *i*^*th*^ row and *j*^*th*^column of a representational dissimilarity matrix (RDM). The bottom right corner depicts a simple activation space, where the meaning of “catfish” is defined by the signal of voxels 1 and 2 in response to the presentation of the “catfish” stimulus. Again, a given metric is used to quantify the dissimilarity 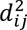 of “catfish” and “snowman”, and a neural RDM is built from the dissimilarities of all combination pairs. Finally, the two RDMs are correlated with Spearman rank correlation (*r*_*s*_) to quantify the representational similarity between the cDSM and neural spaces.

To combine word vectors, scholars developed *compositional* distributional semantics models (cDSMs), namely algebraic functions that take as their argument constituent vectors to generate novel, complex ones 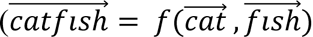; Baroni, Bernardi, et al., 2014; Mitchell & Lapata, 2010). To our knowledge, neuroimaging studies of minimal semantic composition have not made use of (c)DSMs, although they have been used to control for confounds and devise stimuli (Krieger-Redwood et al., 2022; Wang et al., 2020; see also Djokic et al., (2020) for a model-driven study of negation). They have, however, been widely adopted in psycholinguistics (Amenta et al., 2020), where some cDSMs have been proposed as cognitive models of complex word semantics (Marelli et al., 2017; Marelli & Baroni, 2015). Indeed, besides their successful application to NLP (Mitchell & Lapata, 2010; Dinu et al., 2013b), cDSMs have been shown to predict explicit judgments of minimal combinations (Ciapparelli et al., 2025; Günther & Marelli, 2016; 2021; 2022; Vecchi et al., 2017), as well as chronometric behavioral measures related to their processing (Günther & Marelli, 2018, 2020a; 202b). Therefore, by specifying what functions are needed to carry out composition and by relying on rich constituent representations (and their known representational similarity with brain regions in the semantic network), cDSMs constitute promising tools for studying the neural correlates of semantic composition.

More specifically, since different functions can be applied to the same word vectors, cDSMs allow us to specify alternative compositional representations of the same stimuli (complemented by non-compositional ones from DSMs, if the compound is familiar). When cDSMs differ exclusively along the combinatorial dimension, model comparisons can be transparently attributed to the compositional functions, which, if interpretable, can be related to (neuro)cognitive theories of conceptual combination. Model comparison can thus inform researchers not only about which regions are correlated with semantic combination but also what type of combination they likely support. In the present work we take this approach and rely on three cDSMs whose computations are interpretable and can be related to theories of conceptual combination advanced in cognitive neuroscience (section 1.1) and computational psycholinguistics (Baroni, 2013; Marelli et al., 2017; Vecchi et al., 2017). Specifically, we focus on the following models:

- **Additive model** (Mitchell & Lapata, 2010): This model computes the sum of constituent word vectors:

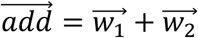

The model can be parsimoniously taken to represent the simple co-activation of constituents’ meanings (i.e., their superimposition) rather than proper composition.
- **Multiplicative model** (Mitchell & Lapata, 2010): This model consists of the component-wise multiplication of a word vector pair, namely:

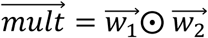

Where ⊙ represents the multiplication of vector components, 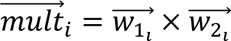. The model can be taken to compute the feature intersection of constituent word vectors (Baroni, 2013), as their semantic components interact to jointly determine the combination’s component (Baron & Osherson, 2011).
- **Compounding as Abstract Operation in Semantic Space (CAOSS) model** (Marelli et al., 2017): The CAOSS model is a multiple linear regression trained to predict the vector of a compound from the vectors of its constituents (see Methods 2.3.2). At inference, the model generates compound representations in two steps. First, constituent words are multiplied by role-specific matrices. Then, the compound representation is obtained by taking the sum of left and right constituent vectors:

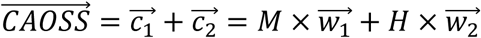

Where the *M* matrix is applied to 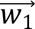 to obtain its role-specific representation as the left constituent 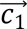 (the same process is applied to the right constituent 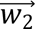 via the *H* matrix). Thus, contrary to the additive and multiplicative models, CAOSS distinguishes words as modifiers or head constituents and assigns those roles in the form of a linear transformation applied to their vector representation.

### 1.3 Putting the pieces together: A model-based re-analysis

In the present work, we re-analyze data aggregated from four fMRI studies, all employing English two-word combinations but varying in task requirements (see Table 1; section 2.1), using the cDSMs described above. Consistent with the definition of semantic composition as a transformation of (semantic) representations, we employ representational similarity analysis (RSA; Kriegeskorte et al., 2008) to compare model and brain representations. Contrary to univariate approaches, RSA does not relate neural activity to a property of the stimulus or task (first-order isomorphism); rather, RSA is concerned with whether neural activation patterns reflect the representational structure of the stimuli (second-order isomorphism), namely the dissimilarity among stimuli in the space of theoretical models (Figure 1; section 2.3). In this way, we set out to test how constituents are combined across the studies considered, testing claims of region-specific computation (section 1.1) by leveraging what we know of model-specific computations (section 1.2). Specifically, we conduct a series of confirmatory RSAs to test the following brain-model correspondences (Figure 2):

- <u>LATL ↔ multiplicative model</u>: It has been claimed that the LATL supports semantic composition via the intersective conjunction of constituent concepts (Coutanche et al., 2018; Baron & Osherson, 2011), which is non-syntactic (Pylkkänen, 2019). Similarly, the multiplicative model operates a form of feature intersection (Baroni, 2013), which is insensitive to the syntactic (morphological) role of constituents (i.e., 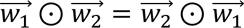). Therefore, we expected to observe a significant representational correspondence between the LATL and the multiplicative model, with the latter offering the best model of LATL representations compared to other theoretical models.
- <u>LAG, RAG ↔ CAOSS model</u>: LAG and RAG are hypothesized to support relational semantic composition based on thematic relations (Boylan et al., 2017; Coutanche et al., 2018; Schwartz et al., 2011). Morphological information (correctly assigning the role of modifier or head to constituents) is essential to build thematic links (e.g., a “house dog” is not a “dog house”). The CAOSS model applies role-specific linear transformations to assign them the typical semantic behavior of modifier or head constituents. Notably, because the modifier and asymmetric 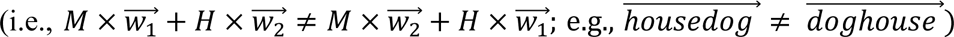. Marelli et al., (2017) demonstrated that CAOSS can predict the effects of relational dominance and relational priming, which depend on the thematic relations connecting constituent meanings (Gagné & Shoben, 1997), while Günther & Marelli, (2022) showed that these are implicitly encoded by the compositional vectors it generates. Therefore, we expect the CAOSS model space to best match the neural space of LAG and RAG.
- <u>LIFG ↔ multiplicative model, CAOSS model</u>: Because specific hypotheses concerning the type of composition supported by the LIFG are currently lacking, we do not advance specific claims of correspondence between cDSMs and this region. However, we include LIFG in our analyses to address the possibility that its involvement in semantic composition might be driven by increased semantic control demands posed by less plausible combinations (Graves et al., 2010; Molinaro et al., 2015; Coutanche et al., 2018). Specifically, we would take a significant model match for either the multiplicative and/or the CAOSS model as evidence for the contrary, that is, of LIFG representing combinatorial information. We also extend these considerations to the AG, for which analogous claims have been advanced (Humphreys et al., 2021; Ralph et al., 2016).

**Figure 2:**
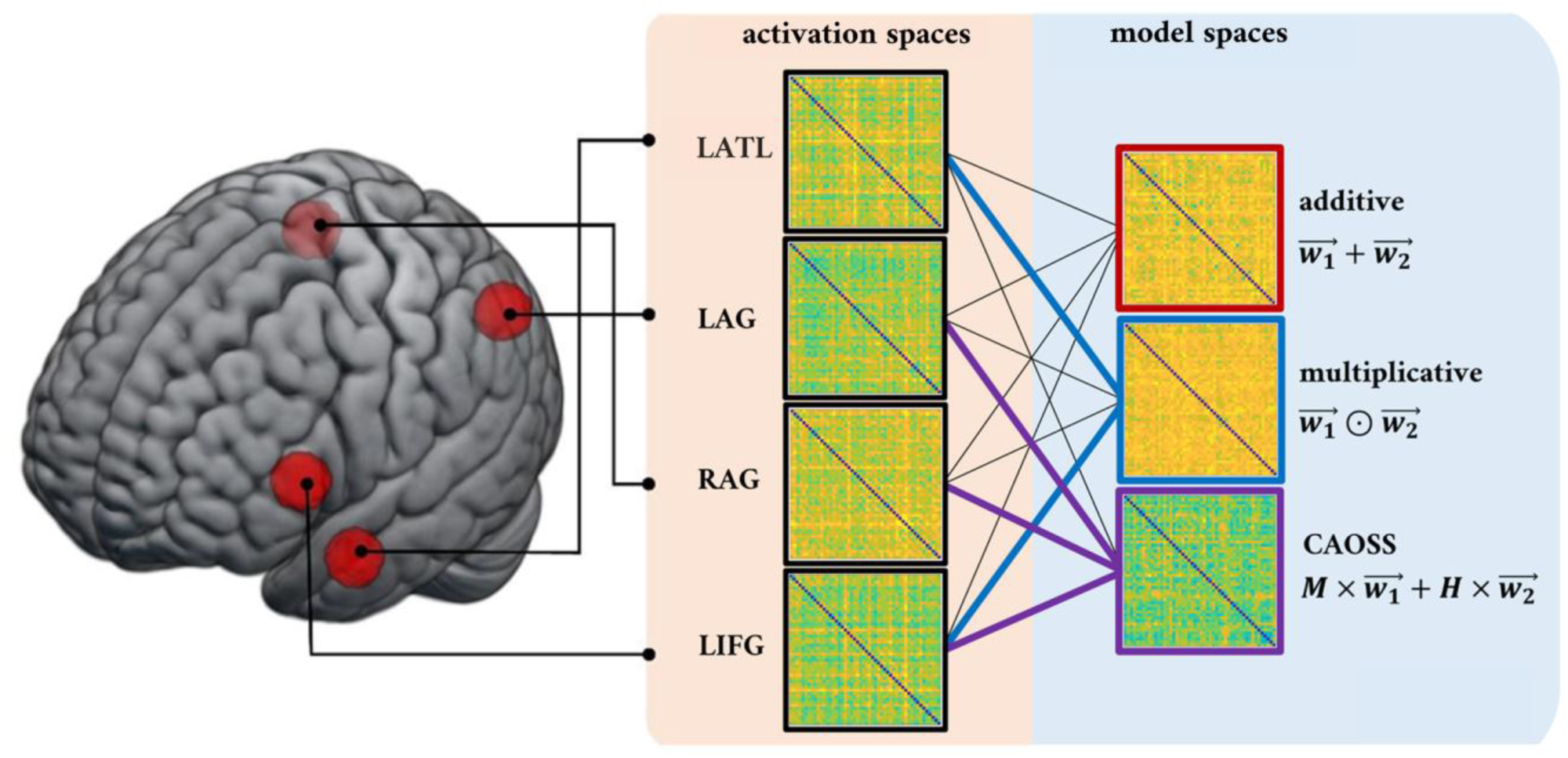
cDSM-ROI correspondences tested in the confirmatory RSA. The RDMs of the four core ROI (spherical ROIs on the left) are correlated with the RDMs of theoretical models. Colored connections indicate correlations of special interest based on claims of specific compositional operations in core ROIs.

**Table 1:** Summary of the stimuli, task, and participant characteristics of the studies included.

| study | stimuli | task | participants |
| --- | --- | --- | --- |
| Graves et al., (2010; experiment 1) | 480 noun-noun phrases, comprising 240 meaningful phrases (e.g., “ski jacket”) and their reverse non-meaningful counterparts (“jacket-ski”). | 1-back task. Participants indicated whether either word of the current phrase matched a word in the same position of the previous phrase (true in 1/6 of trials). | 23 participants (13 females); all were healthy adult native English speakers, had normal or corrected-to-normal vision, and were right-handed. Mean age was 24.2 (SD: 3.0). |
| Graves et al., (2010; experiment 2) | 400 noun-noun phrases; 200 meaningful phrases and their reverse non-meaningful counterparts. | Meaningfulness judgements. Participants indicated whether phrases were meaningful, non- | 22 participants (15 females); all were healthy adult native English speakers, had normal or corrected-to-normal vision, |
|  | 200 nonword phrases were used as baseline trials. | meaningful, or made of “nonwords.” | and were right handed. Mean age was 24.7 (SD: 5.4). |
| Wang et al., (2020) | 312 noun-noun phrases. Of these, 96 were place-thing phrases (e.g., “temple-priest”), 96 were category-exemplar phrases (e.g., “dessert-brownie”), 96 whole-part phrases (e.g., “dog-tail”) and 24 were unrelated (e.g., “mascara-spoon”) | Semantic relation judgements. Participants indicated whether they could think of a relation between the constituents or not. 1/3 of related phrases were primed (i.e., preceded by a phrase with the same relation type) | 18 participants (11 females); all were healthy adult native English speakers, and were right handed. Age ranged from 18 to 28 years. |
| Graves et al., (2022) | 192 adjective-noun phrases, varying along the socialness-figurativeness dimensions (i.e., social figurative, social literal, nonsocial figurative, nonsocial literal) | Familiarity judgements. Participants indicated whether each phrase they were seeing was familiar or unfamiliar. | Subset of 22 neurotypical participants; all were healthy adult native English speakers, had normal or corrected-to-normal vision, and were right handed. Mean age was 21.3. |

In this context, the additive model is used as a baseline model for the simple non-compositional juxtaposition of constituents’ meanings (Baron & Osherson, 2011). Because all core ROIs (except RAG) have been largely implicated in general semantic processing, we expected to find representational similarities between the additive model and all core ROIs, while we also expect those core ROIs to represent compositional information beyond simple addition. Notably, besides LATL, LAG, RAG, and LIFG, other regions from the broader semantic network have been implicated in semantic composition, albeit less consistently (Frankland & Greene, 2020a; Pylkkänen, 2019). Furthermore, few neuroimaging studies have used RSA to study semantic composition (but see Baron & Osherson, 2011; Momenian et al., 2021; Wang et al., 2020), and none with computational models of semantic composition. For this reason, we complement our confirmatory ROI analysis with an exploratory searchlight RSA conducted in the broader semantic network.

Finally, our modeling approach and data allowed us to test two additional hypotheses advanced in psycholinguistic research, namely the claim that semantic composition is attempted automatically, and that this attempt is carried out in parallel with retrieval from long-term memory (Günther & Marelli, 2020; Libben, 2014), compatible with a parallel architecture of compositional processing (Baggio, 2020; 2021) and neuroimaging evidence of automatic semantic processing (Liuzzi et al., 2021):

- <u>Automaticity</u>: If semantic composition was attempted automatically upon the presentation of a suitable stimulus, we should expect to observe combinatorial semantic representations in the brain even when the task does not require explicit semantic processing. To test this claim, we re-run the confirmatory analyses described above using only the data from one study where the task could be accomplished relying on word form alone (study 1; Graves et al., 2010), expecting to nonetheless replicate the results observed when all studies are considered.
- <u>Parallel routes of semantic composition and retrieval</u>: Two of the four studies we re-analyze include familiar stimuli, namely familiar combinations (e.g., “airplane”) for which a non-compositional vector representation is available in the DSM’s semantic space. This allows us to compare, for the same stimuli, non-compositional and compositional representations, dissociating the “storage” (i.e., lexicalized meaning retrieval) and “computation” (i.e., automatic and active semantic composition) routes that are claimed to operate in parallel (Libben, 2014). We focus on this stimulus subset and conduct a second exploratory analysis in the broad semantic network, comparing compositional and non-compositional representations. Following Libben (2014), we expect both composition and retrieval to be attempted; thus, we expect to find representational similarities driven by both compositional and non-compositional models.

## 2. Materials and methods

### 2.1 Experimental materials

We searched for fMRI studies that met the following criteria. First, because combinatorial semantics is expressed differently across languages (e.g., compound productivity, headedness; Libben et al., 2020), we decided to focus on a single language. We chose English due to the larger number of fMRI studies available, its compounding productivity (Libben et al., 2020), the availability of computational resources and their psycholinguistic evaluation (Baroni, Dinu, et al., 2014; Günther & Marelli, 2020, 2022; Marelli et al., 2017; Mitchell & Lapata, 2010). Therefore, we required participants to be native speakers of English. Relatedly, we focused on studies that employed English stimuli, either adjective-noun or noun-noun phrases, requiring both constituents to be presented visually and simultaneously. Finally, to apply the same data analytic procedures across studies, we considered studies for which raw structural and functional data, either DICOM or un-preprocessed NIfTI files, were available. We contacted the authors of studies meeting the above criteria and obtained data from four studies: Graves et al., (2010; experiments 1 and 2), Graves et al., (2022), and Wang et al., (2020). Table 1 displays the main characteristics of the stimuli, tasks, and participants involved in each study.

### 2.2 Imaging methods

Table 2 summarizes the characteristics of fMRI data acquisition across the four studies included. The same preprocessing and data analysis pipelines were applied to all studies, accounting for study-specific differences in image acquisition (i.e., TR and acquisition sequence).

**Table 2:** Summary of the characteristics of image acquisition for the four studies included.

| **study** | **Scanning and trial structure** | **image acquisition** |
| --- | --- | --- |
| Graves et al., (2010; experiment 1) | The scanning session was split into four runs. On each trial, a phrase was displayed for 1 sec and replaced with a fixation cross. For each run, 120 phrase trials were randomly interspersed with 100 fixation trials (mean duration: 3.6 sec, SD: 2.4). | 3 T GE Excite system scanner. Functional volumes were acquired using a gradient-echo echoplanar sequence (matrix = 64×64, 33 interleaved axial slices, voxels dimension 3.5×3.5 mm and slice thickness 3.0 mm with a 0.5 mm gap, TR = 2000 msec, TE = 25 msec). T1 structural images were acquired as a set of 134 contiguous axial slices using a spoiled-gradient-echo sequence. Functional volumes were acquired with gradient-echo echoplanar sequence. |
| Graves et al., (2010; experiment 2) | Identical to Graves et al., (2010; experiment 1). | Identical to Graves et al., (2010; experiment 1) |
| Wang et al., (2020) | The scanning session was split into four runs. On each trial, a phrase was shown for 2 sec, followed by a variable duration fixation cross (ranging from 1 to 6 sec). | 3 T Siemens MRI scanner. Functional volumes were acquired using an EPI sequence (matrix = 64×64, 34 descending axial slices, 3.5 mm isotropic voxels, TR = 2000 msec, TE = 22 msec). T1 structural images were acquired with gradient-echo inversion pulse sequence (1-mm isotropic voxels). |
| Graves et al., (2022) | The scanning session was split into four runs. On each trial, a phrase was displayed for 1.5 sec, followed by a fixation cross for 0.5 sec and a fixation cross for a randomly jittered inter-trial interval (mean duration: 2.5 sec). | 3 T S TIM Trio scanner. Functional volumes acquired as T2*-weighted gradient-echo single-shot EPI (matrix = 64 × 64, 35 interleaved axial slices, 3 mm isotropic voxels, TR = 1900 msec, TE = 2.52 msec). T1 structural MPRAGE images (1 mm isotropic voxels). |

#### 2.2.1 fMRI preprocessing

We carried out fMRI data preprocessing with SPM12 software (Wellcome Trust Centre for Neuroimaging, London, UK). Volumes were spatially realigned, unwarped, slice-time corrected, and co-registered to structural T1 images. We employed the TsDiffAna toolbox (http://www.fil.ion.ucl.ac.uk/spm/ext/#TSDiffAna) to detect volumes of exceedingly high activation that were potentially due to artifacts. Subsequent analyses were carried out in subject space, with the exception of group-level searchlight RSA, for which volumes were normalized to the standard MNI template.

#### 2.2.2 General Linear Model Estimation

We conducted univariate analyses using the general linear model (GLM) approach implemented in SPM12. The primary goal of these analyses was to generate beta maps for each phrase stimulus, which would serve as the foundation for subsequent representational similarity analyses (RSAs). Across all studies, each phrase was presented only once, requiring the estimation of beta maps from single trial presentations. Such estimation, however, poses a significant challenge. Indeed, the combination of trial-by-trial estimation and rapid event-related designs can lead to unreliable beta estimates due to the collinearity arising from the overlap of BOLD signals from adjacent trials, as noted by Mumford et al. (2012). To counter this problem, we implemented a staggered estimation approach. In this approach, we estimated beta coefficients for each trial at intervals of three trials and used another regressor to model all remaining trials falling within the trials of interest. Specifically, we employed three separate GLMs. The first GLM included a predictor for each individual phrase in trials 1, 4, 7, etc., while a single predictor coded for all left-out trials (i.e., trials 2, 3, 5, 6, 8, and so on). The second GLM included predictors for trials 2, 5, 8, and so on, and the third for trials 3, 6, 9, etc., each with their corresponding left-out trials. In these GLMs, predictors coded for the identity of each phrase or group of phrases against an implicit baseline. Notably, we made a deliberate choice to exclude covariates coding for high-level conditions (e.g., whether the word pair was “forward” and meaningful, or “reverse” and non-meaningful; Graves et al., 2010). This decision was based on two considerations. First, since high-level conditions varied across studies, their inclusion could result in beta estimates that depend more on the specific manipulation carried out in each study. This could potentially confound the aggregation of results across different studies. Second, most high-level conditions code for semantic dimensions, which could represent part of the information that should be captured by semantic models. Including these as covariates could inadvertently remove meaningful signals. Additionally, it could hinder model comparison, as semantic models might differ in their sensitivity to the semantic dimensions underlying these high-level conditions.

#### 2.2.3 Representational dissimilarity matrices

After describing each stimulus with a beta map, we compared the activation similarity among stimuli to construct representational dissimilarity matrices (RDMs). First, given a subject and a region of interest (ROI), we constructed an RDM by computing the Mahalanobis distance (a distributional distance metric showing good results for complex stimuli; Bobadilla-Suarez, 2019) between pairs of stimulus phrases, each described by a vector of beta estimates, covering all voxels in the ROI (Nili et al., 2014). From all RDMs, we removed the rows and columns corresponding to stimuli with unreliable beta estimates. First, we used TsDiffAna to detect volumes with anomalous activation. If any of these volumes fell within 2 to 10 sec after the onset of a stimulus, we considered the beta estimates for that stimulus to be unreliable and removed its corresponding row and column from the RDM. Second, single-trial beta estimates of adjacent trials can be unreliable due to collinearity, leading to inflated positive correlations between temporally adjacent trials (Mumford et al., 2014). Because phrases from 1/6 adjacent trials share a constituent in Graves et al., (2010; experiment 1), and phrases from 1/3 adjacent trials share their semantic relation in Wang et al., (2020), the overestimated similarity of adjacent trials might lead to inflated Type I errors (Mumford et al., 2014). Therefore, we removed the first off-diagonal elements from all RDMs.

### 2.3 Theoretical models

We defined three classes of theoretical models: baseline models coding for possible confounding variables, distributional semantics models coding for non-compositional semantic representations, and cDSMs coding for compositional semantic representations. Each model defined a theoretical RDM (or candidate RDM; Nili et al., 2014) used in RSA analyses. Figure 3 displays the second-order dissimilarities among theoretical models.

**Figure 3:**
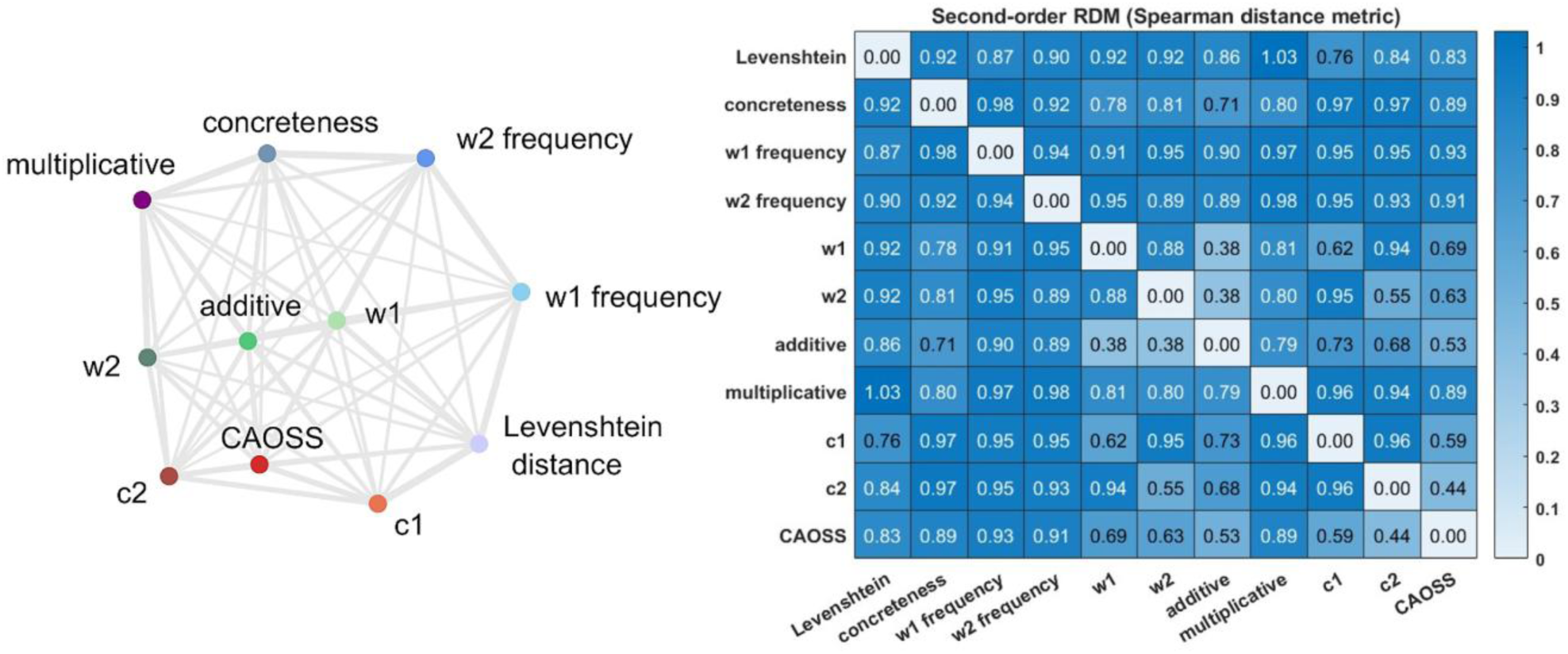
Second-order dissimilarities among theoretical models for all stimuli across all studies (run RDM excluded due to subject-and study-specific construction). (left) Multidimensional scaling based on Spearman distance, computed with Nili et al., (2014) toolbox. (right) Second-order dissimilarity matrix; each cell shows the distance of two theoretical RDMs, computed as 1 – their Spearman correlation.

#### 2.3.1 Baseline models

To better isolate semantics from confounding representations, we controlled semantic models for information coded by the following five theoretical RDMs:

- **Run:** The run RDM coded for whether trials belonged to the same (0) or different (1) functional runs.
- **Orthographic overlap:** The degree of orthographic overlap between two phrases was quantified by their Levenshtein distance, namely the minimum number of single character edits needed to change a phrase into the other. The orthographic overlap RDM was populated by the Levenshtein distances among all pairs of phrases.
- **Left and right constituent frequency:** We quantified word frequency based on the log-transformed word frequency data from the Baroni et al., (2014) corpus, the same corpus used to train the semantic models (section 2.3.2). We constructed a RDM coding for the absolute difference in word frequency of the left constituents from all trial pairs (e.g., the absolute value of the difference between the word frequency of “snow” and “cat” for the “snow man” and “cat fish” trials). We constructed a RDM for the right constituents analogously.
- **Concreteness:** We employed Brysbaert et al., (2014) concreteness norms and defined the concreteness of a word pair to be the average concreteness of its constituents^1^. We constructed a concreteness RDM based on the absolute difference in concreteness between trials.

#### 2.3.2 Distributional semantics models

Word vectors were obtained from the semantic spaces provided by Baroni, Dinu, et al., (2014); specifically, word vectors were derived from a ∼ 2.8 billion word corpus obtained from the concatenation of the web-collected ukWaC (Baroni et al., 2009), a 2008 English Wikipedia dump, and the print-media-based British National Corpus (BNC consortium, 2007). Vectors were computed with the *cbow* version of the *word2vec* model (Mikolov et al., 2013), a prediction-based DSM (Mandera et al., 2017) aimed at predicting a word given its context. The model was trained on the corpus described above, choosing the parameter set producing the best empirical results in Baroni et al., (2014), namely where 400-dimensional vectors were trained to predict words within a context window of five words (negative sampling with k = 10, subsampling with t = 1e−5). These vectors represent the lexicalized meaning of single words and form the basis on which compositional vectors were generated. Based on these vectors, we defined three semantic models:

- **Left constituents**: 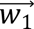. These vectors represented the meaning of the left constituents, i.e., the first word of each phrase;
- **Right constituents**: 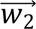 . These vectors represented the meaning of the right constituents, i.e., the second word of each phrase.

We quantified the semantic dissimilarity of two stimuli by computing 1 – the cosine similarity of their vectors. For each semantic model, we constructed a corresponding RDM coding for the semantic dissimilarity of stimuli, resulting in two semantic RDMs: *w*_1_ RDM and *w*_2_ RDM.

#### 2.3.3 Compositional distributional semantics models

From the semantic space described in the previous section, we defined three compositional distributional semantic models:

- **Additive model**: 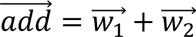 (Mitchell & Lapata, 2010). These vectors represented the unweighted sum of the left and right constituents. Note that, while technically a compositional model, when used as a model of neural representations a more parsimonious interpretation of the vector addition is that it codes for the mere (non-compositional) co-representations of constituents’ features.
- **Multiplicative model**: 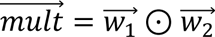 (Mitchell & Lapata, 2010). These vectors represent the elementwise multiplication of constituents’ features.
- **CAOSS model**: 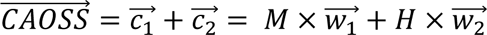 (Marelli et al., 2017). Following Günther & Marelli, (2021), the training data consisted of a set of triplets of (*left constituent, right constituent, compound*) from the union of four noun-noun compound databases: The 629-compound database by Juhasz et al., (2014), the 1,865-compound database by Günther & Marelli, (2018), the 2,861-compound database by Kim et al., (2019), and the 8,376-compound database by Gagné et al., (2019). To guarantee reliable model representations, we kept those for which the compound and its constituents had a raw frequency higher than 50 in the Baroni et al., (2014) corpus. This resulted in a training set of 4,429 triplets. We estimated CAOSS parameters (i.e., the M and H matrices) with the DISSECT toolkit (Dinu et al., 2013a).

We then built *additive, multiplicative* and *CAOSS* RDMs coding for the trials’ semantic dissimilarity as estimated by these models (i.e., 1 – the cosine similarity of additive, multiplicative and CAOSS representations).

### 2.4 Representational similarity analyses

After having defined theoretical and empirical RDMs, we performed the following representational similarity analyses (RSAs).

#### 2.4.1 Confirmatory RSA

We performed RSA in four regions of interest, namely LATL, LAG, RAG, and LIFG. We defined the core ROI as spherical masks with a diameter of 9 voxels, centered on the average activation peaks reported in the literature (Supplementary Table S1). The MNI coordinates we used are: LATL (x = –48.1, y = 0.1, z = –24.5), LAG (x = –40.2, y = –66.6, z = 36.9), RAG (x = 46.9, y = –62.3, z = 35.2), and LIFG (x = –47.4, y = 21.3, z = 7.7). We quantified the second-order similarity between the representational spaces of a given ROI and theoretical model by computing the Spearman correlation between their RDMs (Kriegeskorte et al., 2008; Nili et al., 2014). Importantly, because we wanted to control for the influence of confounding dimensions, we conducted *partial* RSA by performing partial Spearman correlations between RDM pairs controlling for the influence of other RDMs. Specifically, for each ROI, we performed a partial RSA for every candidate semantic (*w*_1_, *w*_2_) and compositional semantic (*additive, multiplicative, CAOSS*) model, controlling for baseline models (i.e., including all baseline models as control variables in partial Spearman correlations). We then conducted statistical inference over the entire sample of participants and studies by computing right-tailed Wilcoxon signed-rank tests, testing whether the partial correlation coefficients were significantly greater than zero (Kriegeskorte et al., 2008; Nili et al., 2014), applying Bonferroni correction for the number of ROIs (i.e., *α* = .05/4). Significant results indicate that a given ROI represents semantic information consistently with a theoretical model and that this second-order similarity cannot be attributed to the confounding dimensions of word frequency, orthographic overlap, concreteness, or run. As stated in section 1.3, we expected to observe a significant representational match between the space of cDSMs and ROIs, with the multiplicative model providing the best representational model of LATL, CAOSS providing the best model of LAG and/or RAG, and either *Multiplicative* or *CAOSS* showing significant representational correspondence with LIFG.

#### 2.4.2 Confirmatory RSA: non-semantic subset

Because evidence suggests that (compositional) semantic processing is automatically enacted (Günther & Marelli, 2020; Liuzzi et al., 2021), we expected to find a representational match for semantic and compositional semantic models even when composition was not required to carry out a task. To test this hypothesis, we focused on Graves et al., (2010) experiment 1, where participants were tasked to indicate if either word in the current phrase matched a word in the same position from the previous phrase. This 1-back task could be accomplished by simply processing surface-level properties of the stimuli, without semantic processing. Therefore, we considered all semantic and compositional models significant in the previous confirmatory analysis (section 2.4.1), and repeated the RSA analyses using only this data subset. We expected to observe the same pattern of model significance in this non-semantic study subset.

#### 2.4.3 Exploratory RSA

Searchlight RSAs were carried out by iteratively considering voxels belonging to a sphere with radius = 3 voxels centered at each voxel across the general semantic network (Jackson, 2021). The analysis was performed in the space of each subject, for every subject and study. For each sphere, the corresponding brain RDM coded for the Mahalanobis distance between pairs of stimuli. Stimuli falling in outlier volumes were discarded. We performed Spearman correlations between brain RDMs and each baseline model, and partial Spearman correlations between brain RDMs and each semantic and compositional semantic model controlling for all baseline models. Searchlights RSAs were implemented with Nili et al., (2014) toolbox. The resulting correlation maps were normalized to reference MNI space, and masked using the intersection of two masks: i) a mask excluding all voxels missing from at least 9 subjects; ii) a mask of the general semantic network peaks from Jackson, (2021) meta-analysis. We conducted group-level analyses over the resulting normalized correlation maps using SPM12, applying a threshold of *α* = .005 at the voxel level, and *α* = .05 at the cluster level, family-wise error (FWE) corrected for multiple comparisons. We did not have specific expectations about where significant clusters could emerge and for which model (with the exception, of course, of the core ROIs).

#### 2.4.4 Exploratory RSA: composition and retrieval

We repeated the searchlight RSAs in the semantic network focusing on the subset of stimuli for which a lexicalized (i.e., non-compositional) vector was available. Specifically, we identified the stimulus phrases for which a vector representation was available in the Baroni et al., (2014) semantic space: 45 stimuli in Graves et al., (2010; experiment 1 and 2), 11 stimuli in Graves et al., (2022), and 4 stimuli in Wang et al., (2020) met this criterion. Due to the limited number of stimuli, we decided to focus on data from Graves et al., (2010; experiment 1 and 2; N = 45). Brain RDMs coded for the Mahalanobis distance between these 45 stimuli (990 pairs), excluding stimuli falling in outlier volumes. Brain RDMs were constructed from voxel spheres (radius = 3 voxels) whose center was systematically varied to cover the general semantic network. In this analysis, theoretical models also included the lexicalized representations of phrase stimuli, 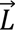, from which we constructed the corresponding *L* RDM. Thus, the semantic representations of a given stimulus phrase (e.g., “apple tree”) were the following: left constituent 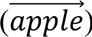, right constituent 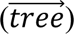, additive model 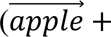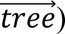, multiplicative model 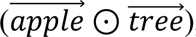, CAOSS model 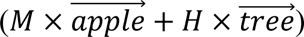, and lexicalized meaning 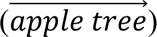. We were especially interested in the unique contribution of the compositional semantic and the lexicalized meaning models. Thus, we conducted partial RSAs controlling for the confounding dimensions of each model. Specifically, given a searchlight voxel sphere, we performed partial Spearman correlations between its brain RDM and the lexicalized meaning RDM, controlling for all baseline and non-compositional semantic models. Similarly, we performed partial Spearman correlations between the brain RDM and each compositional model (i.e., *mult* and *CAOSS*, independently) controlling for baseline models and non-compositional semantic models. The resulting partial correlation maps were normalized to standard MNI space, and masked with the intersection of a mask of the general semantic network and a mask excluding all voxels missing from at least 5 subjects. Once again, we performed group-level analyses over the resulting maps applying an alpha threshold of *α* = .005 at the voxel level, and of *α* = .05 at the cluster level, FWE-corrected for multiple comparisons. In line with the claim that active composition and retrieval processes are simultaneously attempted, we expected to observe significant clusters of compositional semantic models *and* lexicalized representations. Although we did not have *a-priori* localization hypotheses, we expected to observe compositional clusters consistent with results from the previous exploratory searchlight RSA (which included all stimuli from all studies).

## 3. Results

### 3.1 Confirmatory analyses

#### 3.1.1 Confirmatory RSA

Table 3 and Figure 4 report the results of semantic and compositional semantic models. Concerning the former, we find significant results for the additive model in LAG (*Z* = 3.56, *N* = 85, *p* < .001), probably driven, at least in part, by the representational match for the right constituent (*Z* = 2.89, *N* = 85, *p* = .002). For compositional models, we observe significant results for the multiplicative model in LIFG (*Z* = 2.30, *N* = 85, *p* = .011), and a match close to the adjusted alpha threshold (*α* = .0125) for the multiplicative model in LATL (*Z* = 2.09, *N* = 85, *p* = .018).

**Figure 4:**
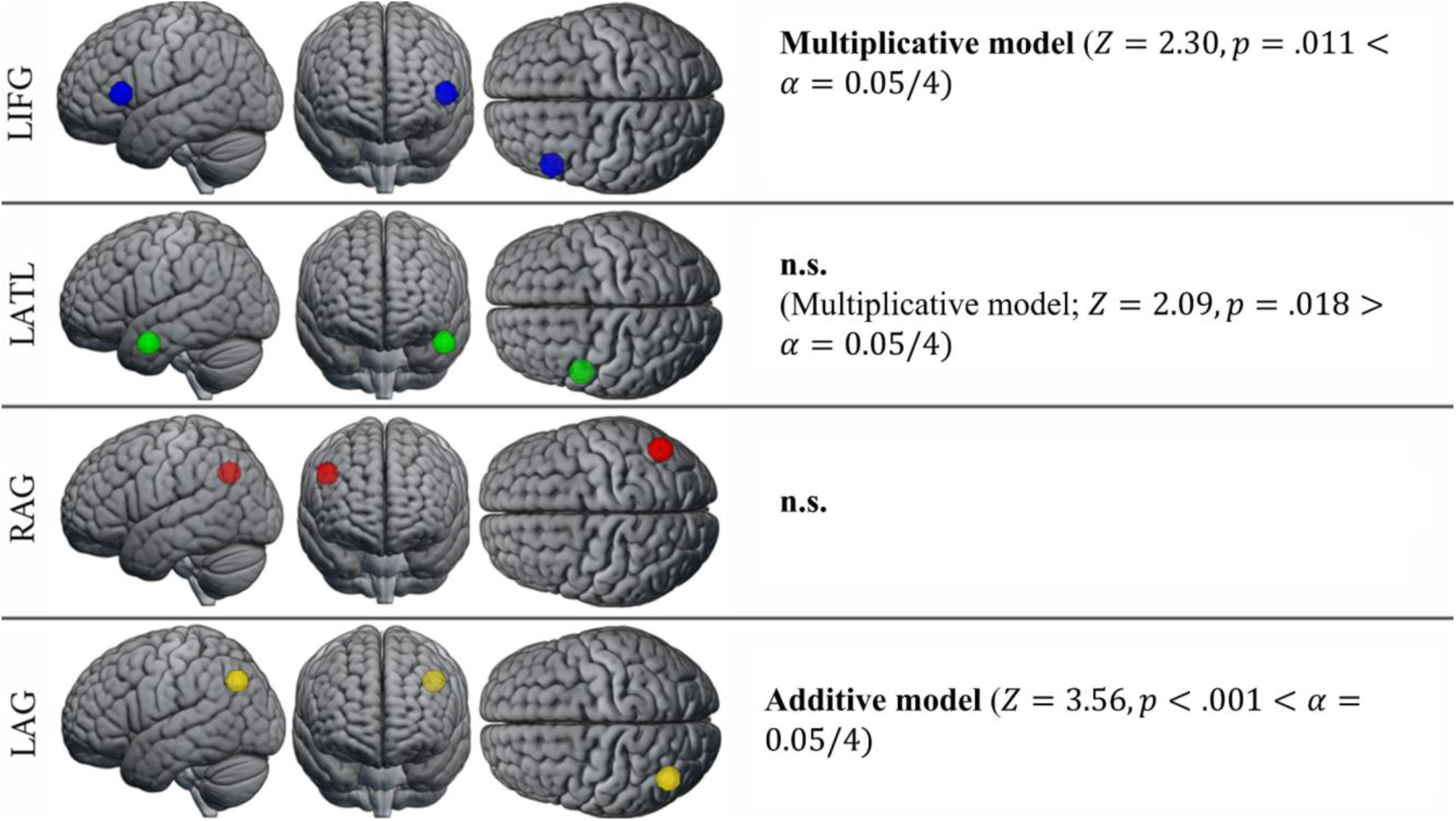
RSA results in the *a-priori-defined* core ROIs.

**Table 3:** Right-tailed Wilcoxon signed-rank test results for semantic and compositional semantic models. Test statistics were computed over the partial Spearman correlation coefficients (i.e., subject-level RSA results) for all subjects and all studies (N = 85). All models were controlled for baseline models. The table reports Z scores and uncorrected *p-*values; results significant after Bonferroni correction (adjusted *α* = .0125) are reported in **bold.** See Supplementary Table S2 for baseline model results.

| RDM predictor | LATL |  | LAG |  | RAG |  | LIFG |  |
| --- | --- | --- | --- | --- | --- | --- | --- | --- |
| | $Z$ | $p$ | $Z$ | $p$ | $Z$ | $p$ | $Z$ | $p$ |
| $w_1$ | 1.50 | .066 | 1.21 | .113 | 0.51 | .306 | 2.14 | .016 |
| $w_2$ | 0.62 | .268 | **2.89** | **.002** | 0.14 | .444 | 1.48 | .069 |
| *add* | 1.39 | .082 | **3.56** | **<.001** | 1.32 | .094 | 2.00 | .023 |
| *mult* | 2.09 | .018 | 1.32 | .093 | 0.54 | .295 | **2.30** | **.011** |
| CAOSS | 0.21 | .418 | 0.50 | .309 | 0.36 | .358 | 0.02 | .493 |

#### 3.1.2 Confirmatory RSA: non-semantic subset

Focusing on the non-semantic task subset of Graves et al., (2010; experiment 1), we repeated RSA for the right constituent and additive model in LAG. Neither the right constituent (*Z* = 0.99, *N* = 23, *p* = .16) nor the additive model (*Z* = 1.14, *N* = 23, *p* = .13) were significant. However, the multiplicative model was still significant in LIFG (*Z* = 2.21, *N* = 23, *p* = .014).

### 3.2 Exploratory analyses

#### 3.2.1 Exploratory RSA

Across all models, a significant representational match was observed only for the multiplicative model. Specifically, searchlight RSA partialling over baseline models revealed two significant clusters, one located in *pars triangularis* of the LIFG (BA45), the other in the left middle superior sulcus (lmSTS; Supplementary Table S3).

#### 3.2.2 Exploratory RSA: composition and retrieval

No significant clusters emerged for any theoretical model^2^. For the lexicalized model, a cluster in the left Fusiform gyrus was close to significance (FWE-corrected cluster *p* = .054; MNI coordinates of peak activation: x = –40, y = –56, z = –14).

**Figure 5:**
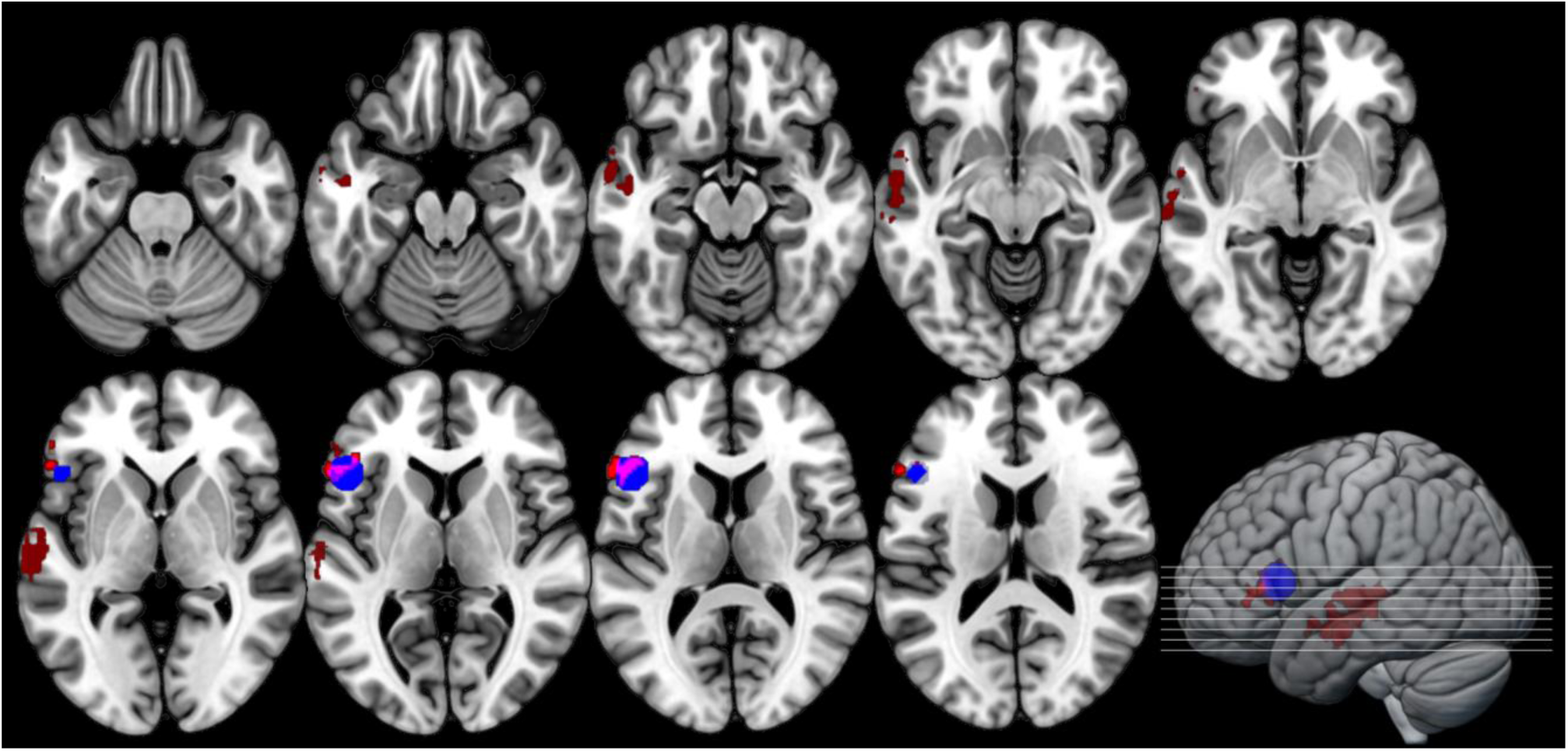
Searchlight RSA results in the broad semantic network. Red: cluster of multiplicative model representations controlling for baseline models; Blue: *a-priori* ROI sphere for LIFG used in confirmatory RSA; Purple: cluster and sphere overlap.

**Figure 6:**
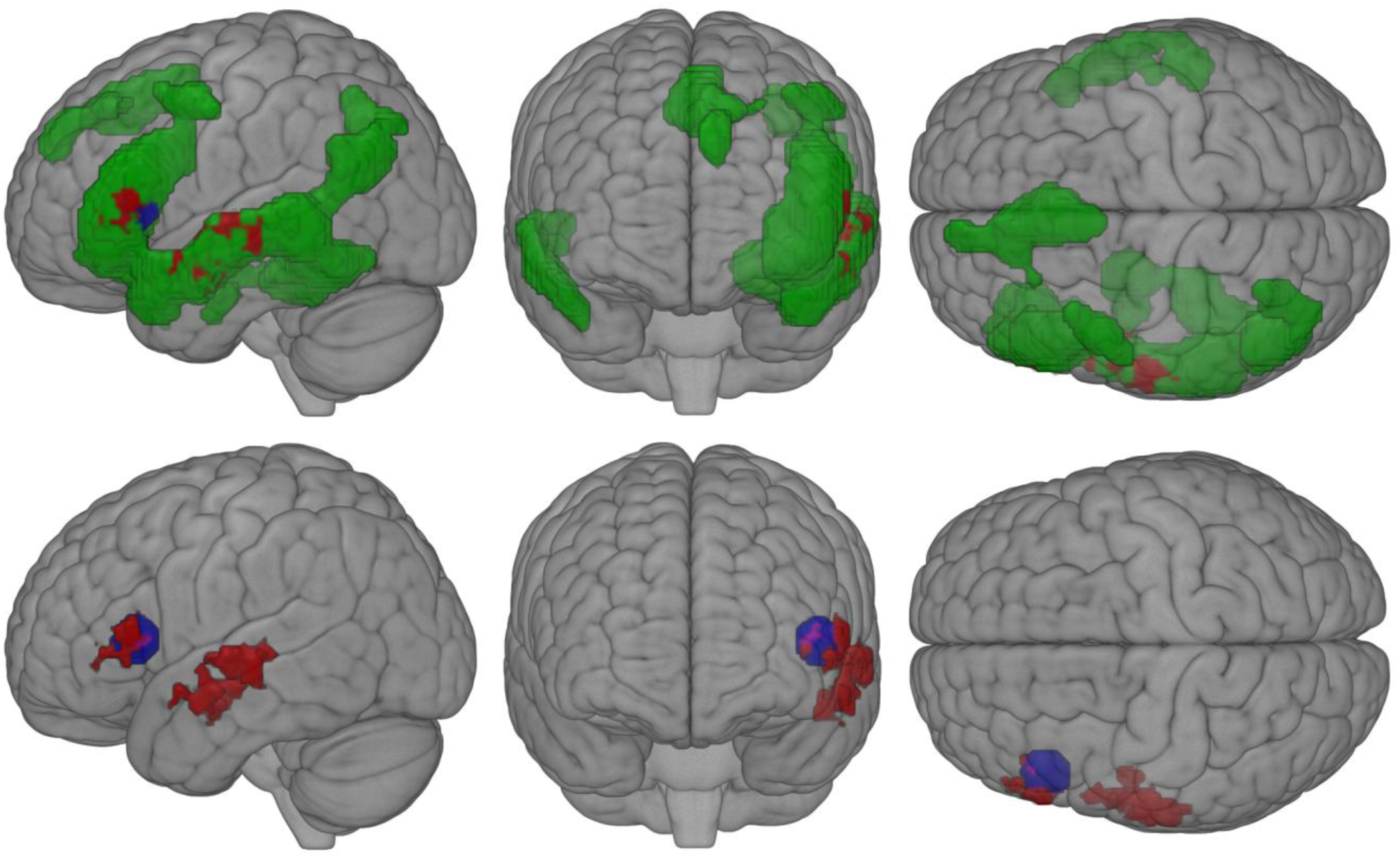
Searchlight RSA results in the broad semantic network. Geen: Mask of the general semantic network (Jackson, 2021). Red: cluster of multiplicative model representations controlling for baseline information; Blue: voxel sphere for LIFG used in the confirmatory RSA. Purple: cluster and sphere overlap.

## 4. Discussion

Semantic composition allows us to construct complex meanings from simpler constituents. Theorists have proposed different ways in which constituents can be combined, some of which have been formalized by computational models such as compositional distributional semantics models (cDSMs). Complementing behavioral studies, neuroimaging can aid the study of semantic composition by tapping into otherwise unobservable representations. Most neuroimaging studies take a univariate approach best suited to identify regions sensitive to high-level contrasts related to semantic composition. However, this approach is more limited in its ability to say *how* composition occurs, that is, what transformations constituent concepts undergo during combination/integration. To address this limitation, the present work asks whether the distributional representations predicted by cDSMs match the fine-grained neural patterns observed in core semantic brain regions. To better target semantic representations beyond specific processing demands, we re-analyzed fMRI data aggregated from four published studies, all employing two-word combinations but differing task requirements. We identified compositional representations in the *pars triangularis* (BA45) of the left inferior frontal gyrus (LIFG), both when aggregating data across all studies and within a subset of studies where the behavioral task did not explicitly require semantic processing. These findings support the hypothesis that the LIFG represents combined concepts and suggest that compositional processing in the LIFG may occur automatically. An exploratory searchlight RSA confirmed these results and revealed an additional cluster of compositional representations in the left middle superior temporal sulcus (lmSTS). Similar, though considerably weaker, representations were also observed in the left anterior temporal lobe (LATL). Unexpectedly, we did not observe compositional representations in the left or right angular gyri (LAG and RAG). We discuss these findings in detail, explore how model characteristics might explain these observations, acknowledge the limitations of the present study, and outline potential avenues for future research.

### 4.1 Compositional representations in the semantic network

*Left Inferior Frontal Gyrus (LIFG):* Compared to LATL and AG, the specific type of semantic composition supported by LIFG is less clear, with theoretical accounts linking LIFG activation to semantic control demands (Graves et al., 2010; Molinaro et al., 2015; Coutanche et al., 2018). Hence, we did not put forward specific hypotheses about which cDSMs would best capture LIFG representations. Instead, we tested whether at least one cDSM could provide a significant model of LIFG, considering a positive result as evidence that LIFG represents semantic compositions beyond specific task demands. This inference is made credible by the diverse array of tasks covered by our re-analysis (Table 1). Indeed, while combinatorial processing could be confounded with semantic control on specific tasks, leading to spurious representational similarities driven by task demands, the same pattern is less likely to drive similarities across multiple, different tasks. Thus, because we find significant representational similarities between the multiplicative model and the LIFG (significant after controlling non-semantic confounds), we conclude that the LIFG represents combinatorial information beyond semantic control demands. Crucially, the searchlight RSA identifies a significant cluster for the multiplicative model in LIFG. This cluster partially overlaps with the spherical mask from the confirmatory RSA analysis, and both are located in LIFG *pars triangularis* (BA45). Moreover, we do not observe non-combinatorial semantic representations in LIFG. This is surprising, as composition depends on constituents’ meaning. We speculate that, if LIFG is rapidly and automatically engaged in composition, the time resolution might be too coarse to identify representations of constituent word meanings before combining them. Indeed, we find multiplicative representations even when considering the 1-back data subset from Graves et al., (2010; experiment 1). These results suggest that LIFG automatically enacts combinatorial processing even when semantic access is not required to solve the task. Consistent with this finding, Liuzzi et al., (2021) showed that LIFG *pars triangularis* was one of three areas where the representations of semantic category information were similar across tasks requiring passive and active semantic access. Semantic category representations, while more pronounced in the active semantic task, were recoverable with a decoder trained on one task and modality and test on the other. Overall, we take the above considerations to support the notion that LIFG *pars triangularis* represents combinatorial semantic information of the multiplicative kind – interpretable as a feature conjunction (Baron & Osherson, 2011; Baroni, 2013) – which is computed automatically and is unlikely to depend on specific semantic control demands.

*Left Anterior Temporal Lobe (LATL):* Confirmatory RSA showed no compositional model was significant in LATL, with the multiplicative model being non-significant after correcting for multiple comparisons (*p* = .072 after Bonferroni correction). While weak, we contend that this result is notable in light of the signal dropout in ATL (Visser et al., 2010), which was not controlled for with appropriate sampling strategies in the fMRI studies we re-analyzed, possibly compounding with the already noisy beta estimates (see Methods 2.2.2). This finding is in line with the hypothesis that the LATL specifically supports composition based on the intersective conjunction of constituent meaning, that is, meaning defined by the properties satisfied by both constituents simultaneously (Coutanche et al., 2018; Poortman & Pylkkänen, 2016), operationalized by the component-wise multiplication of semantic vectors (Baroni, 2013). Indeed, closest to our approach, Baron & Osherson, (2011) sampled the multivoxel BOLD patterns in response to images of faces differing in gender and age, targeting both constituent properties (i.e., “male”, “female”, “adult”, “child”) and their combinations (“woman”, “man”, “girl”, and “boy”) demonstrating that, in LATL and posterior cingulate cortex alone, the activation patterns of combinations were predicted by the multiplication of the activation patterns of their constituents. On the other hand, besides not reaching significance after correction, there are reasons to question this result. First, besides multiplicative representations, Baron & Osherson, (2011) find that additive representations can be recovered from the LATL (and many other regions; see also Baron et al., 2010). Instead, we do not find significant representational similarities between the LATL and the additive model, nor between the LATL and each constituent individually. This goes against the vast evidence of LATL’s centrality in semantic processing more generally and in semantic integration in particular (Ralph et al., 2016), which we hypothesized should lead to representing constituent meaning and their addition, besides multiplication. Thus, overall, while consistent with previous findings of combinatorial processing of the intersective kind in LATL and with our expectations of ROI-model correspondence, this result awaits to be confirmed in future studies.

*Left and Right Angular Gyrus (LAG and RAG):* Contrary to our hypotheses, we did not observe compositional representations in the angular gyri (AG). This finding aligns with the theoretical account proposed by Humphreys et al. (2021), which argues that the AG may not represent semantic information per se. The inconsistent involvement of the AG in semantic cognition studies may be attributable to task complexity acting as a confounding factor in contrasts such as word versus nonword or combinatorial versus non-combinatorial stimuli. Specifically, given the involvement of the left angular gyrus (LAG) in the default mode network (DMN), task complexity could drive the observed deactivation of the AG in univariate studies of semantic composition (see also Ralph et al., 2016). Nevertheless, it remains possible that the AG does support semantic composition, and our compositional DSMs (cDSMs) simply did not provide the appropriate compositional function to detect it. In this regard, Frankland and Greene (2020) link the AG to the DMN, suggesting that the DMN – rather than serving as a repository of semantic knowledge – supports conceptual combination by providing multimodal, context-dependent, and knowledge-based information needed to infer appropriate relationships among constituents. These relationships are frequently complex and challenging to systematize. On this account, cDSMs, as a class of models, might be too simple and rigid to explain combination in the DMN, and consequently in AG^3^ (e.g., they are based on decontextualized word representations, and on textual information alone). Interestingly, however, we find highly significant representational similarities between the additive model and LAG (likely driven, at least in part, by the right constituent), but not RAG. This finding suggests that at least the left (Binder et al., 2009; Jackson, 2021) AG is involved in semantic representation in contrast to the Humphreys et al. (2021) account discussed above. Thus, overall, our findings are consistent with the claim that the reported involvement of AG in conceptual combination might be due to task demands (Humphreys et al., 2021), but also support the notion that LAG does represent semantic information, as widely reported (Binder et al., 2009; Jackson, 2021).

*General semantic network*: Exploratory analyses identify significant clusters of multiplicative representations in the middle portion of the superior temporal sulcus (lmSTS) and in *pars triangularis* of the LIFG (BA45). Extensive evidence has linked LIFG to semantic memory (Binder et al., 2009; Jackson, 2021; Ralph et al., 2016), including more specific evidence of representational similarities between this region and DSMs (Carota et al., 2017; Liuzzi et al., 2020; Zhang et al., 2020). Extensive is also the evidence for the involvement of STS in language processing (Humphries et al., 2006; Pallier et al., 2011; Malik-Moraleda et al., 2022). Looking more specifically at composition, however, previous works implicated STS in processing thematic relations and, more specifically, agent-patient relations (i.e., who did what to whom; Frankland & Greene, 2020a, 2020b), which is incompatible with the multiplicative model’s symmetry. While these results are challenging to reconcile with existing evidence for the localization of semantic composition, we note that the latter are overwhelmingly based on univariate analyses, the complementation of which motivated the exploratory analyses we are discussing. A degree of divergence between the former and the latter is, in this sense, expected, and might inform future studies.

### 4.2 Modeling considerations

Notably, across both confirmatory and exploratory analyses, the multiplicative model consistently emerged as the only cDSM and, in the exploratory RSA, the only theoretical model displaying significant representational similarities with target brain regions. Because cDSMs were tested against data from diverse study settings, which share only the most fundamental properties, we speculate that the good performance of the multiplicative model stems from its simplicity. The fact that we could observe multiplicative representations in the 1-back subset corroborates this interpretation. Indeed, if multiplicative combination is automatically enacted regardless of the need to access or compute semantic information, it naturally follows that one should be able to observe multiplicative representations in other task settings, assuming the same stimulus type (adjective-noun and noun-noun phrases). Nonetheless, similarities with a simple alternative such as the additive model were not observed as pervasively and consistently as those of the multiplicative model. Therefore, the superior performance of the multiplicative model cannot be reduced only to simplicity.

The CAOSS model was never significant, despite predicting compound processing and representation across different behavioral measures and languages (Günther, Marelli, et al., 2020; Günther & Marelli, 2018, 2020, 2022; Marelli et al., 2017). Mirroring the considerations above, CAOSS might be too complex to generate semantic representations consistently present across tasks and thus preserved in their aggregate. Alternatively, while CAOSS predicts a diverse array of behavioral results related to compound processing, the representations that drive such predictions may just not be good models of the neural representations that underlie those behaviors, suggesting that more is needed to develop a cDSM that can account for both.

### 4.3 Limitations and future directions

By considering data coming from diverse fMRI studies, we were able to conduct analyses at a scale larger than usual experiments, posing a stringent test to the generalizability of computational representations across contexts. However, a major downside stemming from this approach is that none of the studies included was optimized for multivariate analyses targeting word-level representations. Indeed, stimuli were presented only once in each study, leading to noisy beta estimates, and BOLD acquisition was not optimized for LATL signal dropout. Studies tailored to our research questions and methodology would have, of course, more statistical power to detect the effects of interest, a limitation that we tried to mitigate with a larger sample size. In this sense, a reduced sample size (55 subjects from Graves et al., (2010); experiment 1 and 2) might be why we fail to observe significant clusters in the searchlight RSA targeting lexicalized and compositional representations, together with the limited amount of stimuli per participant.

Second, our methodology suffers from one key limitation of DSMs, namely their lack of grounding. Theoretical accounts of conceptual combination postulate a central role of sensorimotor information, which is claimed to drive combination in dynamic interaction with linguistic information (Lynott & Connell, 2010), while brain regions implicated in semantic composition are connected to modality-specific regions (e.g., the “hub-and-spoke” model of ATL; Lambon Ralph et al., 2010; Ralph et al., 2016). Indeed, Günther, Petilli, et al., (2020) demonstrate that a CAOSS model trained on vision vectors explains compound word processing times over and above its DSM-based counterpart. Therefore, future studies might benefit from considering semantic representations enriched with extralinguistic information.

Finally, as we discussed in the introduction, we attempted to shift the focus of semantic composition studies from an “activation” perspective, relating univariate responses to high-level contrasts that involve combination, to a “representational” one, where semantic composition is defined as a systematic transformation of meaning-carrying representations. Three key ingredients characterize this framework: the representations of constituent concepts, the representation of their combined meaning and, arguably the end goal of the research endeavor, the transformation that connects the two. Reverberi et al., (2012) provide an example of this approach in the domain of rule representation. Specifically, they obtained the multivoxel BOLD representations of conditional rules and their conjunctive combination and showed that decoders trained on constituent rule identity could be used to recover the identity of their combination successfully. From this perspective, an ingredient missing from the present study is the neural representation of constituent information. Future studies might collect the neural representation of constituent units besides those of their combination; cDSMs could then directly operate on the former to predict the latter (indeed, see Baron et al., 2010; Baron & Osherson, 2011).

## 5. Conclusions

The present work explored the neural bases of semantic composition by testing the representational similarity of combinatorial stimuli described by activation patterns in core regions of interest and in the theoretical spaces defined by compositional distributional semantics models. To better target semantic representations beyond specific processing demands, we reanalyzed fMRI data aggregated from four published studies differing in task requirements. We find evidence for (non-compositional) semantic representations in the left angular gyrus. Converging evidence indicates the use of multiplicative combinatorial representations in the *pars triangularis* of the left inferior frontal gyrus (BA45), suggesting that the latter represents combinatorial information – of the “intersective conjunction” or “feature intersection” kind (Coutanche et al., 2018; Poortman & Pylkkänen, 2016) – beyond task demands and even when semantic processing is not explicitly required. Overall, our model-driven approach clarifies which brain regions represent combinatorial information across contexts and processing demands, offering evidence for some specific operations that may underlie semantic composition in these areas.

## Acknowledgments

We are grateful to Silvia Bunge for sharing her data with us and to Katherine Alfred for assisting in the data-sharing procedure. The contribution of Marco Marelli was supported by the European Union (ERC-COG-2022, BraveNewWord, 101087053). Views and opinions expressed are however those of the authors only and do not necessarily reflect those of the European Union or the European Research Council Executive Agency. Neither the European Union nor the granting authority can be held responsible for them.

## Contributions

**MC:** Conceptualization, Data curation, Formal analysis, Methodology, Software, Visualization, Writing - original draft; **MM:** Methodology, Funding acquisition, Writing - review & editing; **WG:** Data curation, Writing - review & editing; **CR:** Methodology, Software, Resources, Writing - review & editing, Supervision.

## Code availability

The code used in this study is openly available at: https://osf.io/3dnqg/?view_only=7df0e90f3a0745549d7563827eb9e49b

## Footnotes

1 Concreteness ratings were missing for 6 first words and 18 second words. In these cases, only the rating of the available constituent

2 In order to make results comparable with the main exploratory RSA (section 2.4.3), we also performed a searchlight RSA for the compositional models controlling only for the baseline models (but not for the non-compositional semantic models). Again, no significant clusters emerged for either the multiplicative or the CAOSS models.

3 We note that CAOSS, compared to the additive and multiplicative models, allows an additional degree of flexibility. Indeed, the M and H matrices interact with the vectors differently (i.e., the same matrix can change a vector much more than another). But even CAOSS has a degree of rigidity, as M and H matrices do not model the interactions between left and right constituents (e.g., the meaning of “cat” in “cat fish” and “cat sitter” will receive an identical transformation by M).

## Supplementary materials

**Table S1:**
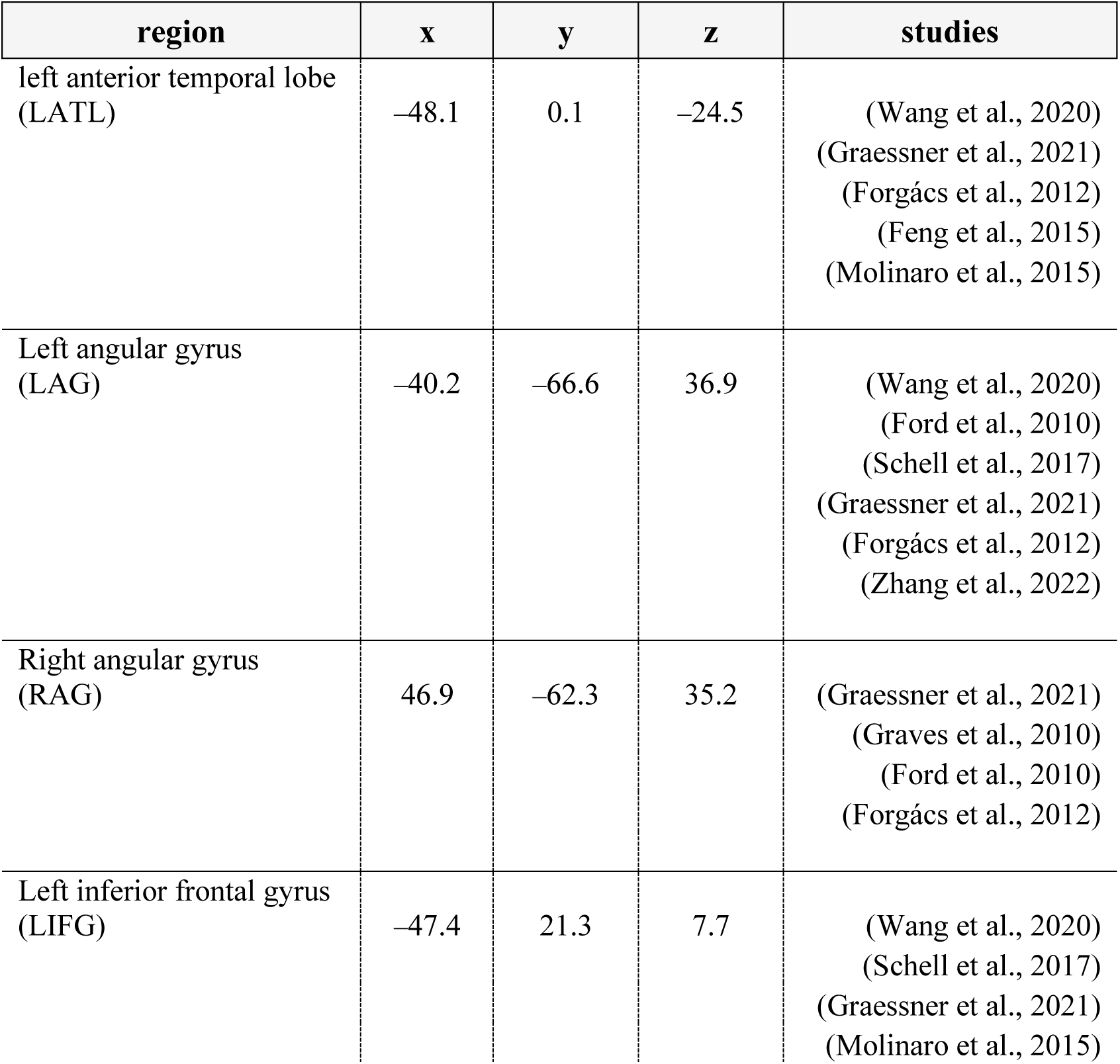
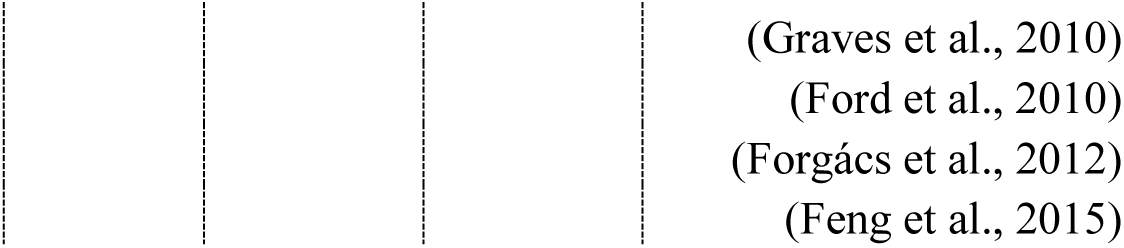
Core ROI coordinates. For each core ROI, the table reports the mean MNI coordinates computed based on the peak activations reported in relevant studies (listed in the rightmost column).

| region | x | y | z | studies |
| --- | --- | --- | --- | --- |
| left anterior temporal lobe (LATL) | −48.1 | 0.1 | −24.5 | (Wang et al., 2020)<br>(Graessner et al., 2021)<br>(Forgács et al., 2012)<br>(Feng et al., 2015)<br>(Molinaro et al., 2015) |
| Left angular gyrus (LAG) | −40.2 | −66.6 | 36.9 | (Wang et al., 2020)<br>(Ford et al., 2010)<br>(Schell et al., 2017)<br>(Graessner et al., 2021)<br>(Forgács et al., 2012)<br>(Zhang et al., 2022) |
| Right angular gyrus (RAG) | 46.9 | −62.3 | 35.2 | (Graessner et al., 2021)<br>(Graves et al., 2010)<br>(Ford et al., 2010)<br>(Forgács et al., 2012) |
| Left inferior frontal gyrus (LIFG) | −47.4 | 21.3 | 7.7 | (Wang et al., 2020)<br>(Schell et al., 2017)<br>(Graessner et al., 2021)<br>(Molinaro et al., 2015) |

**Table S2:** Core ROI results. Right-tailed Wilcoxon signed-rank test results for all models. Test statistics were computed over the Spearman correlation coefficients (i.e., subject-level RSA results) for all subjects and all studies (N = 85). RSAs were performed with Spearman correlations; partial RSAs controlling for confounding models were performed with partial Spearman correlations (models controlled in such a manner are specified with the *p* subscript). The table reports Z scores and uncorrected *p-*values; results significant after Bonferroni correction (adjusted *α* = .0125) are reported in **bold.**

| RDM predictor | LATL |  | LAG |  | RAG |  | LIFG |  |
| --- | --- | --- | --- | --- | --- | --- | --- | --- |
| | $Z$ | $p$ | $Z$ | $p$ | $Z$ | $p$ | $Z$ | $p$ |
| *run* | **7.186** | **<.001** | **7.808** | **<.001** | **7.554** | **<.001** | **7.195** | **<.001** |
| *Levenshtein distance* | -0.272 | .607 | -0.911 | .819 | 0.438 | .331 | -0.311 | .622 |
| *w<sub>1</sub> frequency* | -0.583 | .720 | -0.074 | .530 | -0.245 | .597 | -0.193 | .576 |
| *w<sub>2</sub> frequency* | -0.692 | .756 | 0.232 | .408 | 1.726 | .042 | -0.596 | .724 |
| *concreteness* | 0.355 | .361 | 0.079 | .469 | -0.197 | .578 | 0.241 | .405 |
| *w<sub>1p</sub>* | 1.503 | .066 | 1.209 | .113 | 0.508 | .306 | *2.138* | *.016* |
| *w<sub>2p</sub>* | 0.618 | .268 | **2.892** | **.002** | 0.140 | .444 | 1.481 | .069 |
| *additive<sub>p</sub>* | 1.393 | .082 | **3.558** | **<.001** | 1.315 | .094 | *1.998* | *.023* |
| *multiplicative<sub>p</sub>* | *2.090* | *.018* | 1.323 | .093 | 0.539 | .295 | **2.300** | **.011** |
| *CAOSS<sub>p</sub>* | 0.206 | .418 | 0.500 | .309 | 0.364 | .358 | 0.018 | .493 |

**Table S3:** Peak activations for the searchlight RSA in the semantic network. Coordinates [x, y, z] in Montreal Neurological Institute (MNI) space and selection of cluster maxima according to SPM12. Searchlight RSA was performed with partial Spearman correlations controlling semantic and compositional semantic models for baseline models.

| model | Cluster $p$<br>(FWE) | Cluster<br>size | Peak Z | MNI | | |
| --- | --- | --- | --- | --- | --- | --- |
| *multiplicative* | .0004 | 452 | 3.74 | -64 | -12 | 2 |
|  |  |  | 3.60 | -60 | -22 | 2 |
|  |  |  | 3.42 | -66 | -24 | -8 |
|  | .030 | 186 | 3.55 | -58 | 24 | 8 |
|  |  |  | 3.47 | -44 | 26 | 8 |
|  |  |  | 3.20 | -52 | 26 | 0 |

## References

Amenta, S., Günther, F., & Marelli, M. (2020). A (distributional) semantic perspective on the processing of morphologically complex words. The Mental Lexicon, 15(1), 62–78. 10.1075/ml.00014.ame

Baggio, G. (2020). Meaning in the Brain I: Relational Semantics. Meaning in the Brain. 10.7551/mitpress/11265.003.0003

Baggio, G. (2021). Compositionality in a parallel architecture for language processing. Cognitive Science, 45(5), e12949. 10.1111/cogs.12949

Baron, S. G., & Osherson, D. (2011). Evidence for conceptual combination in the left anterior temporal lobe. NeuroImage, 55(4), 1847–1852. 10.1016/j.neuroimage.2011.01.066

Baron, S. G., Thompson-Schill, S. L., Weber, M., & Osherson, D. (2010). An early stage of conceptual combination: Superimposition of constituent concepts in left anterolateral temporal lobe. Cognitive Neuroscience, 1(1), 44–51. 10.1080/17588920903548751

Baroni, M. (2013). Composition in distributional semantics. Linguistics and Language Compass, 7(10), 511–522. 10.1111/lnc3.12050

Baroni, M., Bernardi, R., & Zamparelli, R. (2014). Frege in Space: A Program of Compositional Distributional Semantics. Linguistic Issues in Language Technology, 9(6), 242–346. https://aclanthology.org/2014.lilt-9.5/

Baroni, M., Bernardini, S., Ferraresi, A., & Zanchetta, E. (2009). The waCky wide web: A collection of very large linguistically processed web-crawled corpora. Language Resources and Evaluation, 43(3), 209–226. 10.1007/s10579-009-9081-4

Baroni, M., Dinu, G., & Kruszewski, G. (2014). Don’t count, predict! A systematic comparison of context-counting vs. context-predicting semantic vectors. 52nd Annual Meeting of the Association for Computational Linguistics, ACL 2014 - Proceedings of the Conference, 1, 238–247. 10.3115/v1/p14-1023

Binder, J. R., Desai, R. H., Graves, W. W., & Conant, L. L. (2009). Where is the semantic system? A critical review and meta-analysis of 120 functional neuroimaging studies. *Cerebral Cortex*, *19*(12), 2767–2796. 10.1093/cercor/bhp055

Bobadilla-Suarez, S., Ahlheim, C., Mehrotra, A., Panos, A., & Love, B. C. (2020). Measures of neural similarity. Computational brain & behavior, 3, 369–383. 10.1007/s42113-019-00068-5

Boylan, C., Trueswell, J. C., & Thompson-Schill, S. L. (2017). Relational vs. attributive interpretation of nominal compounds differentially engages angular gyrus and anterior temporal lobe. Brain and Language, 169, 8–21. 10.1016/j.bandl.2017.01.008

Brysbaert, M., Warriner, A. B., & Kuperman, V. (2014). Concreteness ratings for 40 thousand generally known English word lemmas. Behavior Research Methods, 46(3), 904–911. 10.3758/s13428-013-0403-5

Carota, F., Kriegeskorte, N., Nili, H., & Pulvermüller, F. (2017). Representational similarity mapping of distributional semantics in left inferior frontal, middle temporal, and motor cortex. Cerebral Cortex, 27(1), 294–309. 10.1093/cercor/bhw379

Caucheteux, C., & King, J. R. (2022). Brains and algorithms partially converge in natural language processing. Communications Biology, 5(1). 10.1038/s42003-022-03036-1

Ciapparelli, M., Zarbo, C., & Marelli, M. (2025). Conceptual Combination in Large Language Models: Uncovering Implicit Relational Interpretations in Compound Words With Contextualized Word Embeddings. Cognitive Science, 49(3), e70048. 10.1111/cogs.70048

Coutanche, M. N., Solomon, S. H., & Thompson-Schill, S. L. (2018). Conceptual Combination. The Big Book of Concepts, 1–22. 10.7551/mitpress/1602.003.0012

Coutanche, M. N., & Thompson-Schill, S. L. (2015). Creating concepts from converging features in human cortex. Cerebral cortex, 25(9), 2584–2593. 10.1093/cercor/bhu057

Dinu, G., & Baroni, M. (2013, August). Dissect-distributional semantics composition toolkit. In Proceedings of the 51st Annual Meeting of the Association for Computational Linguistics: System Demonstrations (pp. 31-36).

Dinu, G., Pham, N. T., & Baroni, M. (2013b). General estimation and evaluation of compositional distributional semantic models. Proceedings of the Workshop on Continuous Vector Space Models and Their Compositionality, 50–58.

Djokic, V. G., Maillard, J., Bulat, L., & Shutova, E. (2020). Modeling affirmative and negated action processing in the brain with lexical and compositional semantic models. ACL 2019 - 57th Annual Meeting of the Association for Computational Linguistics, Proceedings of the Conference, 5155–5165. 10.18653/v1/p19-1508

Malik-Moraleda, S., Ayyash, D., Gallée, J., Affourtit, J., Hoffmann, M., Mineroff, Z., … & Fedorenko, E. (2022). An investigation across 45 languages and 12 language families reveals a universal language network. Nature neuroscience, 25(8), 1014–1019. 10.1038/s41593-022-01114-5

Feng, G., Chen, Q., Zhu, Z., & Wang, S. (2015). Separate brain circuits support integrative and semantic priming in the human language system. Cerebral Cortex, 26(7), 3169–3182. 10.1093/cercor/bhv148

Ford, J. H., Verfaellie, M., & Giovanello, K. S. (2010). Neural correlates of familiarity-based associative retrieval. Neuropsychologia, 48(10), 3019–3025. 10.1016/j.neuropsychologia.2010.06.010

Forgács, B., Bohrn, I., Baudewig, J., Hofmann, M. J., Pléh, C., & Jacobs, A. M. (2012). Neural correlates of combinatorial semantic processing of literal and figurative noun noun compound words. NeuroImage, 63(3), 1432–1442. 10.1016/j.neuroimage.2012.07.029

Frankland, S. M., & Greene, J. D. (2020a). Concepts and Compositionality: In Search of the Brain’s Language of Thought. Annual Review of Psychology, 71, 273–303. 10.1146/annurev-psych-122216-011829

Frankland, S. M., & Greene, J. D. (2020b). Two Ways to Build a Thought: Distinct Forms of Compositional Semantic Representation across Brain Regions. Cerebral Cortex, 30(6), 3838– 3855. 10.1093/cercor/bhaa001

Gagné, C. L., & Shoben, E. J. (1997). Influence of thematic relations on the comprehension of modifier-noun combinations. Journal of Experimental Psychology: Learning Memory and Cognition, 23(1), 71–87. 10.1037/0278-7393.23.1.71

Gagné, C. L., Spalding, T. L., & Schmidtke, D. (2019). LADEC: The Large Database of English Compounds. Behavior Research Methods, 51(5), 2152–2179. 10.3758/s13428-019-01282-6

Graessner, A., Zaccarella, E., Friederici, A. D., Obrig, H., & Hartwigsen, G. (2021). Dissociable contributions of frontal and temporal brain regions to basic semantic composition. Brain Communications, 3(2). 10.1093/braincomms/fcab090

Graessner, A., Zaccarella, E., & Hartwigsen, G. (2021). Differential contributions of left-hemispheric language regions to basic semantic composition. Brain Structure and Function, 226(2), 501–518. 10.1007/s00429-020-02196-2

Graves, W. W., Binder, J. R., Desai, R. H., Conant, L. L., & Seidenberg, M. S. (2010). Neural correlates of implicit and explicit combinatorial semantic processing. NeuroImage, 53(2), 638– 646. 10.1016/j.neuroimage.2010.06.055

Graves, W. W., Levinson, H., Coulanges, L., Cahalan, S., Cruz, D., Sancimino, C., Bal, V. H., & Rosenberg-Lee, M. (2022). Neural differences in social and figurative language processing on the autism spectrum. Neuropsychologia, 171(April), 108240. 10.1016/j.neuropsychologia.2022.108240

Günther, F., & Marelli, M. (2016). Understanding karma police: The perceived plausibility of noun compounds as predicted by distributional models of semantic representation. PLoS ONE, 11(10), 1–36. 10.1371/journal.pone.0163200

Günther, F., & Marelli, M. (2018). Enter Sandman: Compound Processing and Semantic Transparency in a Compositional Perspective. Journal of Experimental Psychology: Learning Memory and Cognition, 45(10), 1872–1882. 10.1037/xlm0000677

Günther, F., & Marelli, M. (2020). Trying to make it work: Compositional effects in the processing of compound “nonwords.” Quarterly Journal of Experimental Psychology (2006), *73*(7), 1082–1091. 10.1177/1747021820902019

Günther, F., & Marelli, M. (2021). CAOSS and transcendence : Modeling role-dependent constituent meanings in compounds. Morphology. 10.1007/s11525-021-09386-6

Günther, F., & Marelli, M. (2022). Patterns in CAOSS: Distributed representations predict variation in relational interpretations for familiar and novel compound words. Cognitive Psychology, 134, 101471. 10.1016/j.cogpsych.2022.101471

Günther, F., Marelli, M., & Bölte, J. (2020). Semantic transparency effects in German compounds: A large dataset and multiple-task investigation. Behavior Research Methods, 52(3), 1208– 1224. 10.3758/s13428-019-01311-4

Günther, F., Petilli, M. A., & Marelli, M. (2020). Semantic transparency is not invisibility: A computational model of perceptually-grounded conceptual combination in word processing. Journal of Memory and Language, 112(July 2019), 104104. 10.1016/j.jml.2020.104104

Günther, F., Rinaldi, L., & Marelli, M. (2019). Vector-Space Models of Semantic Representation From a Cognitive Perspective: A Discussion of Common Misconceptions. Perspectives on Psychological Science, 14(6), 1006–1033. 10.1177/1745691619861372

Harris, Z. S. (1954). Distributional Structure. WORD, 10(2–3), 146–162. 10.1080/00437956.1954.11659520

Humphreys, G. F., Lambon Ralph, M. A., & Simons, J. S. (2021). A Unifying Account of Angular Gyrus Contributions to Episodic and Semantic Cognition. Trends in Neurosciences, 44(6), 452–463. 10.1016/j.tins.2021.01.006

Humphries, C., Binder, J. R., Medler, D. A., & Liebenthal, E. (2006). Syntactic and semantic modulation of neural activity during auditory sentence comprehension. Journal of cognitive neuroscience, 18(4), 665–679. 10.1162/jocn.2006.18.4.665

Jackson, R. L. (2021). The neural correlates of semantic control revisited. NeuroImage, 224(July 2020), 117444. 10.1016/j.neuroimage.2020.117444

Juhasz, B. J., Lai, Y. H., & Woodcock, M. L. (2014). A database of 629 English compound words: ratings of familiarity, lexeme meaning dominance, semantic transparency, age of acquisition, imageability, and sensory experience. Behavior Research Methods, 47(4), 1004–1019. 10.3758/s13428-014-0523-6

Kenett, Y. N., & Thompson-Schill, S. L. (2020). Novel conceptual combination can dynamically reconfigure semantic memory networks. 10.31234/osf.io/crp47

Kim, S. Y., Yap, M. J., & Goh, W. D. (2019). The role of semantic transparency in visual word recognition of compound words: A megastudy approach. Behavior Research Methods, 51(6), 2722–2732. 10.3758/s13428-018-1143-3

Krieger-Redwood, K., Steward, A., Gao, Z., Wang, X., Halai, A., Smallwood, J., & Jefferies, E. (2022). Creativity in verbal associations is linked to semantic control. Cerebral Cortex, 1–13. 10.1093/cercor/bhac405

Kriegeskorte, N., Mur, M., & Bandettini, P. (2008). Representational similarity analysis - connecting the branches of systems neuroscience. Frontiers in Systems Neuroscience, 2(NOV), 1–28. 10.3389/neuro.06.004.2008

Kumar, A. A. (2020). Semantic memory: A review of methods, models, and current challenges. In *Psychonomic Bulletin and Review*. Psychonomic Bulletin & Review. 10.3758/s13423-020-01792-x

Lambon Ralph, M. A., Sage, K., Jones, R. W., & Mayberry, E. J. (2010). Coherent concepts are computed in the anterior temporal lobes. Proceedings of the National Academy of Sciences of the United States of America, 107(6), 2717–2722. 10.1073/pnas.0907307107

Landauer, T. K., & Dumais, S. T. (1997). A Solution to Plato’s Problem: The Latent Semantic Analysis Theory of Acquisition, Induction, and Representation of Knowledge. Psychological Review, 104(2), 211–240. 10.1037/0033-295X.104.2.211

Libben, G. (2014). The nature of compounds: A psychocentric perspective. Cognitive Neuropsychology, 31(1–2), 8–25. 10.1080/02643294.2013.874994

Libben, G., Gagné, C. L., & Dressler, W. U. (2020). The Representation and Processing of Compound Words. Word Knowledge and Word Usage, 336. 10.1093/acprof:oso/9780199228911.001.0001

Liuzzi, A. G., Aglinskas, A., & Fairhall, S. L. (2020). General and feature-based semantic representations in the semantic network. Scientific Reports, 10(1), 1–12. 10.1038/s41598-020-65906-0

Liuzzi, G. A., Ubaldi, S., & Laurence, S. (2021). Representations of conceptual information during automatic and active semantic access. Neuropsychologia, 160(July), 107953. 10.1016/j.neuropsychologia.2021.107953

Lynott, D., & Connell, L. (2010). Embodied conceptual combination. Frontiers in Psychology, 1(NOV), 1–14. 10.3389/fpsyg.2010.00212

Mack, M. L., Preston, A. R., & Love, B. C. (2013). Decoding the brain’s algorithm for categorization from its neural implementation. Current Biology, 23(20), 2023–2027. 10.1016/j.cub.2013.08.035

Mandera, P., Keuleers, E., & Brysbaert, M. (2017). Explaining human performance in psycholinguistic tasks with models of semantic similarity based on prediction and counting: A review and empirical validation. Journal of Memory and Language, 92, 57–78. 10.1016/j.jml.2016.04.001

Marelli, M., & Baroni, M. (2015). Affixation in semantic space: Modeling morpheme meanings with compositional distributional semantics. Psychological Review, 122(3), 485–515. 10.1037/a0039267

Marelli, M., Gagné, C. L., & Spalding, T. L. (2017). Compounding as Abstract Operation in Semantic Space: Investigating relational effects through a large-scale, data-driven computational model. Cognition, 166, 207–224. 10.1016/j.cognition.2017.05.026

Martin, A. E., & Baggio, G. (2020). Modelling meaning composition from formalism to mechanism. Philosophical Transactions of the Royal Society B: Biological Sciences, 375(1791), 1–7. 10.1098/rstb.2019.0298

Mikolov, T. (2013). Distributed Representations ofWords and Phrases and their Compositionality. EMNLP 2016-Conference on Empirical Methods in Natural Language Processing, Proceedings, 1389–1399. 10.18653/v1/d16-1146

Mikolov, T., Chen, K., Corrado, G., & Dean, J. (2013). Efficient estimation of word representations in vector space. 1st International Conference on Learning Representations, ICLR 2013 - Workshop Track Proceedings, 1–12. 10.48550/arXiv.1301.3781

Mitchell, J., & Lapata, M. (2010). Composition in Distributional Models of Semantics. Cognitive Science, 34(8), 1388–1429. 10.1111/j.1551-6709.2010.01106.x

Mitchell, T. M., Shinkareva, S. V., Carlson, A., Chang, K. M., Malave, V. L., Mason, R. A., & Just, M. A. (2008). Predicting human brain activity associated with the meanings of nouns. science, 320(5880), 1191–1195. 10.1126/science.1152876

Molinaro, N., Paz-Alonso, P. M., Duñabeitia, J. A., & Carreiras, M. (2015). Combinatorial semantics strengthens angular-anterior temporal coupling. Cortex, 65, 113–127. 10.1016/j.cortex.2015.01.004

Momenian, M., Radman, N., Rafipoor, H., Barzegar, M., & Weekes, B. (2021). Compound words are decomposed regardless of semantic transparency and grammatical class: An fMRI study in Persian. Lingua, 259. 10.1016/j.lingua.2021.103120

Mumford, J. A., Davis, T., & Poldrack, R. A. (2014). The impact of study design on pattern estimation for single-trial multivariate pattern analysis. NeuroImage, 103, 130–138. 10.1016/j.neuroimage.2014.09.026

Mumford, J. A., Turner, B. O., Ashby, F. G., & Poldrack, R. A. (2012). Deconvolving BOLD activation in event-related designs for multivoxel pattern classification analyses. NeuroImage, 59(3), 2636–2643. 10.1016/j.neuroimage.2011.08.076

Nili, H., Wingfield, C., Walther, A., Su, L., Marslen-Wilson, W., & Kriegeskorte, N. (2014). A Toolbox for Representational Similarity Analysis. PLoS Computational Biology, 10(4). 10.1371/journal.pcbi.1003553

Pallier, C., Devauchelle, A. D., & Dehaene, S. (2011). Cortical representation of the constituent structure of sentences. Proceedings of the National Academy of Sciences, 108(6), 2522–2527. 10.1073/pnas.1018711108

Parrish, A., & Pylkkänen, L. (2022). Conceptual Combination in the LATL With and Without Syntactic Composition. Neurobiology of Language, 3(1), 46–66. 10.1162/nol_a_00048

Pereira, F., Lou, B., Pritchett, B., Ritter, S., Gershman, S. J., Kanwisher, N., Botvinick, M., & Fedorenko, E. (2018). Toward a universal decoder of linguistic meaning from brain activation. Nature Communications, 9(1). 10.1038/s41467-018-03068-4

Poortman, E. B., & Pylkkänen, L. (2016). Adjective conjunction as a window into the LATL’s contribution to conceptual combination. Brain and Language, 160, 50–60. 10.1016/j.bandl.2016.07.006

Price, Amy R., Bonner, M. F., Peelle, J. E., & Grossman, M. (2015). Converging evidence for the neuroanatomic basis of combinatorial semantics in the angular gyrus. Journal of Neuroscience, 35(7), 3276–3284. 10.1523/JNEUROSCI.3446-14.2015

Price, Amy Rose, Peelle, J. E., Bonner, M. F., Grossman, M., & Hamilton, R. H. (2016). Causal evidence for a mechanism of semantic integration in the angular Gyrus as revealed by high-definition transcranial direct current stimulation. Journal of Neuroscience, 36(13), 3829–3838. 10.1523/JNEUROSCI.3120-15.2016

Pylkkänen, L. (2019). The neural basis of combinatory syntax and semantics. Science, 366(6461), 62–66. 10.1126/science.aax0050

Ralph, M. A. L., Jefferies, E., Patterson, K., & Rogers, T. T. (2016). The neural and computational bases of semantic cognition. Nature Reviews Neuroscience, 18(1), 42–55. 10.1038/nrn.2016.150

Reverberi, C., Görgen, K., & Haynes, J. D. (2012). Compositionality of rule representations in human prefrontal cortex. Cerebral Cortex, 22(6), 1237–1246. 10.1093/cercor/bhr200

Schell, M., Friederici, A. D., & Zaccarella, E. (2022). Neural classification maps for distinct word combinations in Broca’s area. Frontiers in Human Neuroscience, 16, 930849. 10.3389/fnhum.2022.930849

Schell, M., Zaccarella, E., & Friederici, A. D. (2017). Differential cortical contribution of syntax and semantics: An fMRI study on two-word phrasal processing. Cortex, 96, 105–120. 10.1016/j.cortex.2017.09.002

Schmidtke, D., Gagné, C. L., Kuperman, V., Spalding, T. L., & Tucker, B. V. (2018). Conceptual relations compete during auditory and visual compound word recognition. Language, cognition and neuroscience, 33(7), 923–942. 10.1080/23273798.2018.1437192

Schrimpf, M., Blank, I. A., Tuckute, G., Kauf, C., Hosseini, E. A., Kanwisher, N., Tenenbaum, J. B., & Fedorenko, E. (2021). The neural architecture of language: Integrative modeling converges on predictive processing. Proceedings of the National Academy of Sciences of the United States of America, 118(45). 10.1073/pnas.2105646118

Schwartz, M. F., Kimberg, D. Y., Walker, G. M., Brecher, A., Faseyitan, O. K., Dell, G. S., Mirman, D., & Coslett, H. B. (2011). Neuroanatomical dissociation for taxonomic and thematic knowledge in the human brain. 10.1073/pnas.1014935108

Solomon, S. H., & Thompson-Schill, S. L. (2020). Feature Uncertainty Predicts Behavioral and Neural Responses to Combined Concepts. Journal of Neuroscience, 40(25), 4900–4912. 10.1523/JNEUROSCI.2926-19.2020

Vecchi, E. M., Marelli, M., Zamparelli, R., & Baroni, M. (2017). Spicy Adjectives and Nominal Donkeys: Capturing Semantic Deviance Using Compositionality in Distributional Spaces. Cognitive Science, 41(1), 102–136. 10.1111/cogs.12330

Visser, M., Embleton, K. V, Jefferies, E., Parker, G. J., & Ralph, M. A. L. (2010). The inferior, anterior temporal lobes and semantic memory clarified: Novel evidence from distortion-corrected fMRI. 10.1016/j.neuropsychologia.2010.02.016

Wang, W. C., Hsieh, L. T., Swamy, G., & Bunge, S. A. (2021). Transient neural activation of abstract relations on an incidental analogy task. Journal of Cognitive Neuroscience, 33(1), 77–88. 10.1162/jocn_a_01622

Westerlund, M., & Pylkkänen, L. (2014). The role of the left anterior temporal lobe in semantic composition vs. semantic memory. Neuropsychologia, 57(1), 59–70. 10.1016/j.neuropsychologia.2014.03.001

Wisniewski, E. J. (1996). Construal and similarity in conceptual combination. Journal of Memory and Language, 35(3), 434–453. 10.1006/jmla.1996.0024

Zhang, L., & Pylkkänen, L. (2015). NeuroImage The interplay of composition and concept speci fi city in the left anterior temporal lobe : An MEG study. NeuroImage, 111, 228–240. 10.1016/j.neuroimage.2015.02.028

Zhang, W., Xiang, M., & Wang, S. (2022). The role of left angular gyrus in the representation of linguistic composition relations. Human Brain Mapping, 43(7), 2204–2217. 10.1002/hbm.25781

Zhang, Y., Han, K., Worth, R., & Liu, Z. (2020). Connecting concepts in the brain by mapping cortical representations of semantic relations. Nature Communications, 11(1), 1–13. 10.1038/s41467-020-15804-w

Ziegler, J., & Pylkkänen, L. (2016). Neuropsychologia Scalar adjectives and the temporal unfolding of semantic composition : An MEG investigation. Neuropsychologia, 89, 161–171. 10.1016/j.neuropsychologia.2016.06.010

